# Single-Cell Multiomics Reveals Distinct Cell States at the Top of the Human Hematopoietic Hierarchy

**DOI:** 10.1101/2021.04.01.437998

**Authors:** Mikael N.E. Sommarin, Parashar Dhapola, Fatemeh Safi, Rebecca Warfvinge, Linda Geironson Ulfsson, Eva Erlandsson, Anna Konturek-Ciesla, Ram Krishna Thakur, Charlotta Böiers, David Bryder, Göran Karlsson

**Affiliations:** Division of Molecular Hematology, Lund Stem Cell Center, Lund University, Sweden

**Keywords:** ScRNA-seq, ScATAC-seq, CITE-seq, HSC, Hematopoiesis, Differentiation, Enhancers, Ageing, Stem cells

## Abstract

The advent of single cell (Sc) genomics has challenged the dogma of haematopoiesis as a tree-like structure of stepwise lineage commitment through distinct and increasingly restricted progenitor populations. Instead, analysis of ScRNA-seq has proposed that the earliest events in human hematopoietic stem cell (HSC) differentiation are characterized by only subtle molecular changes, with hematopoietic stem and progenitor cells (HSPCs) existing as a continuum of low-primed cell-states that gradually transition into a specific lineage (CLOUD-HSPCs). Here, we combine ScRNA-seq, ScATAC-seq and cell surface proteomics to dissect the heterogeneity of CLOUD-HSPCs at different stages of human life. Within CLOUD-HSPCs, pseudotime ordering of both mRNA and chromatin data revealed a bifurcation of megakaryocyte/erythroid and lympho/myeloid trajectories immediately downstream a subpopulation with an HSC-specific enhancer signature. Importantly, both HSCs and lineage-restricted progenitor populations could be prospectively isolated based on correlation of their molecular signatures with CD35 and CD11A expression, respectively. Moreover, we describe the changes that occur in this heterogeneity as hematopoiesis develops from neonatal to aged bone marrow, including an increase of HSCs and depletion of lympho-myeloid biased MPPs. Thus, this study dissects the heterogeneity of human CLOUD-HSPCs revealing distinct HSPC-states of relevance in homeostatic settings such as ageing.

## Introduction

HSCs are essential for maintaining hematopoietic homeostasis and have substantial clinical relevance as the critical cellular component of stem cell transplantations. Hematopoiesis is also the conceptual paradigm to which other somatic stem cell systems are most frequently compared and which has its foundation in the plethora of studies that have attempted to structure murine hematopoiesis[1]. High-purity prospective isolation of human HSCs was long hampered by the absence of permissive xenograft models. However, the development of improved immune-deficient mouse strains, e.g. NOD/LtSz-*scid*/Il*2rg*^-/-^ (NSG) mice[2] has paved the way for the definition of human HSCs as Lin^-^CD34^+^CD38^-^CD45RA^-^CD90^+^CD49f^+^ cells (CD49F^+^HSC)[3]. Yet, 80-90% of cells within this HSC population are not bona fide stem cells as measured in xenograft models. Instead, subclasses of cells exist within populations currently defined as HSCs that exhibit variable reconstitution kinetics, lineage output, and self-renewal potential. In addition, the markers used currently to define human HSCs have mainly been established using cord blood (CB)[3] which might not allow direct translation to other ontogenetic stages due to changes in HSC function over time[4, 5]. For instance, it has been proposed that HSCs have a lymphoid-biased differentiation potential early in life that shifts into a myeloid-bias with age [6-8] and which is paralleled by increases of immunophenotypic HSCs that have decreased reconstitution potential[6, 7, 9].

Immunophenotypic characterization along with evaluation of lineage output using functional assays has long been the mainstay to define HSC heterogeneity and lineage priming[3, 10]. Simultaneously, hematopoietic differentiation has been viewed as a marked hierarchical process where lineage-restriction is gained in a stepwise manner from HSCs through distinct progenitor populations. Recent technological advancements allow for investigation of the transcriptome[11, 12], as well as the epigenome[13, 14] at single-cell (Sc) resolution using ScRNA-sequencing (ScRNAseq) and single cell Assay for Transposase-Accessible Chromatin (ScATACseq), respectively. Initial ScRNA-seq studies described the early stages of hematopoietic differentiation as a continuum of low-primed undifferentiated hematopoietic stem- and progenitor cells (CLOUD-HSPCs) that gradually acquire transcriptomic lineage priming and lose self-renewal capacity[15, 16].

Due to the continuous nature of lineage differentiation, several bioinformatic methods have been developed to quantify the lineage relations of cells[17, 18]. Generally, these methods place the cells in a trajectory and order them along a pseudotime to investigate dynamic changes in gene expression causing lineage determination. A recent study using inheritable transcribed barcodes and ScRNA-seq showed that methods such as these readily captures the lineage relationships of cells. Additionally, it was shown that ScRNA-seq cannot fully capture the complexities of hematopoietic lineage commitment[19].

In contrast, for cell type-classification the epigenome is superior to the transcriptome and ATAC-seq analysis improves cluster-purity compared to RNA-seq[20]. Interestingly, cell type-specification can be further amended by separately analyzing the chromatin accessibility within distal elements as compared to promoters. Distal elements encompass several different genetic features, including enhancers known to be highly cell type-specific and critical early regulators of transcriptional alterations during cellular differentiation[21]. These features make enhancer accessibility a promising tool to define molecularly and functionally distinct subpopulations in heterogeneous fractions of closely related cells.

Utilizing ScATAC-seq and pseudotime analysis we here define the dynamic changes in enhancer accessibility and transcription factor activity that occur as lineage commitment is initiated within the CLOUD-HSPCs. By integrating ScATAC-seq with CITE-seq (Cellular indexing transcriptome and epitome by sequencing)[22] we are able to link enhancer phenotypes to gene expression signature and epitome. Importantly, we demonstrate the existence of lineage-biased cell states within CLOUD-HSPCs, and prospectively isolate lympho/myeloid restricted MPPs based on the cell-surface marker CD11A. Moreover, within the immunophenotypically defined HSCs, we identify a subpopulation of cells that contain a primitive HSC-specific enhancer signature correlating with multilineage reconstitution capacity as well as cell-surface expression of CD35. This CD35^+^ HSC population is molecularly conserved in umbilical cord blood as well as ageing bone marrow. However, distinct age-related effects on heterogeneity are observed where lineage-biased MPPs are altered in relative representation as well as gene expression. Together, our results dissect CLOUD-HSPC heterogeneity into distinct isolable cell types and reveal the effects on HSPC heterogeneity associated with ageing.

## Results

### Characterizing human HSPC heterogeneity using CITE-seq analysis

The current model for human hematopoiesis, including the definition of cell-surface markers for human HSC and progenitor purification has been established in human cord blood (CB), while the translation to other ontogenetic stages is less well defined[3]. Changes in HSPC heterogeneity and function with age are well established[6, 23]. For example, although the frequencies of immunophenotypic HSCs increase with age, engraftment capacity is decreased[6]. Additionally, HSPCs from young individuals have a lymphoid-biased differentiation potential while older HSCs are myeloid-biased[6]. To dissect the heterogeneity of HSPCs at different stages of life, we applied a strategy to directly combine single-cell gene expression analysis with high-throughput immunophenotypic profiling (CITE-Seq) on neonatal, young and aged HSPC populations (Figure1A). First, to identify cell surface markers that might separate the human HSC population, CD49f^+^HSCs from umbilical cord blood (CB) mononuclear cells were analysed against a panel of 342 different antibodies that recognize individual cell surface markers (FigureS1A). Individual validation resulted in the selection of 9 cell surface markers with the potential to reveal heterogeneity. These included CD11A, a previously defined negative marker for HSCs in mouse [24] and CD105 (ENG), a marker of human HSCs in fetal liver [25].

To capture age-associated alterations in heterogeneity within the human HSC population, we compared the results from our immunophenotypic screens of CB to our previously published data on aged BM (aBM, >55 years of age)[26]. Within the HSC-enriched Lin^-^CD34^+^CD38^-^ (CD38^-^HSCs) population, we could define 20 markers that shift in expression in aBM (Figure1B), including the HSC marker CD90 that has previously been shown to increase with age[6], as well as CD41 that has been suggested to mark age-related changes of the hematopoietic hierarchy in mouse[23]. Together, 43 cell surface markers potentially relevant for HSPC heterogeneity were compiled for inclusion in subsequent single-cell molecular analyses.

**Figure 1.**
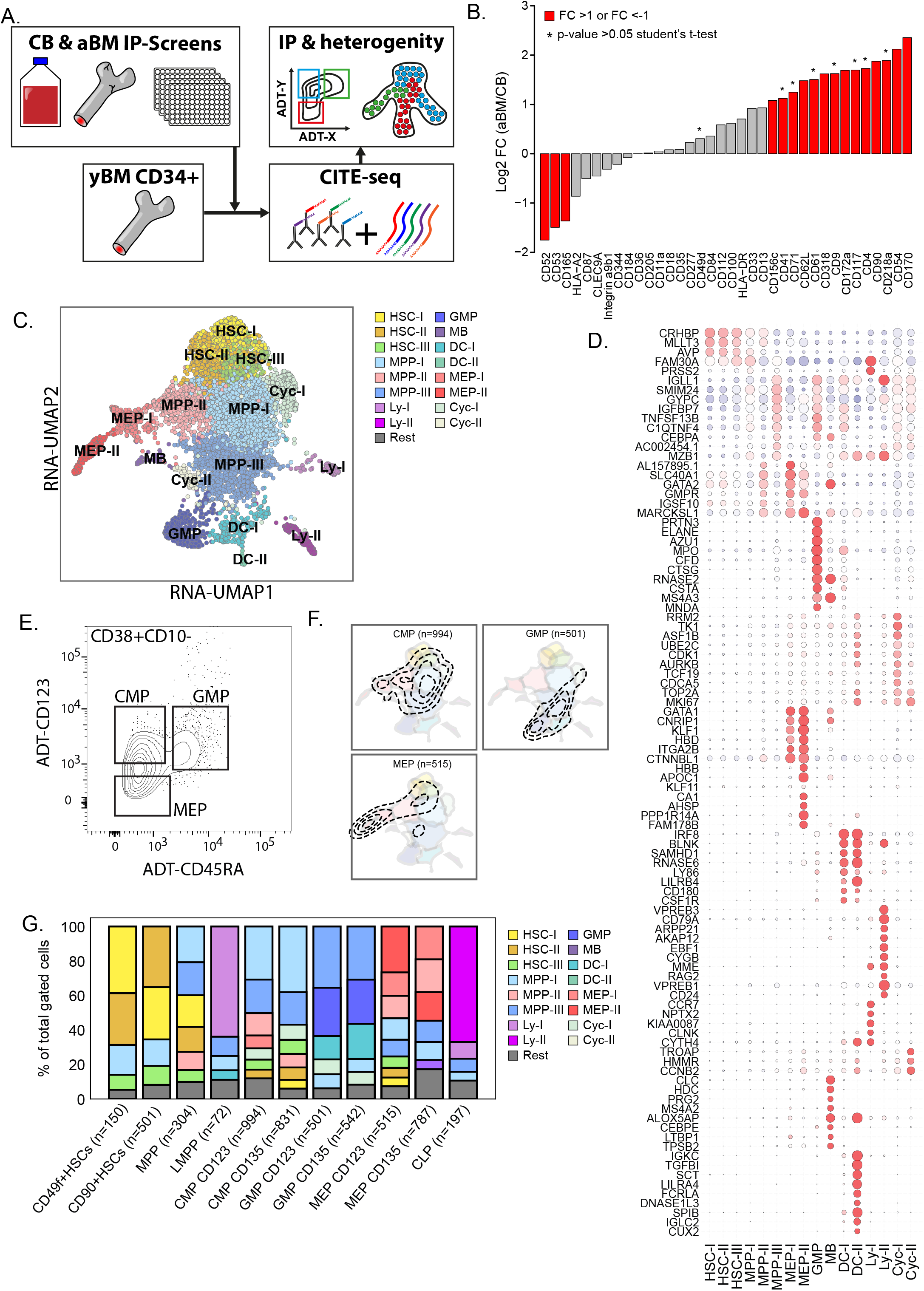
Characterization of immunophenotypic- and transcriptomic heterogeneity of CD34+HSPCs. A. Schematic of experimental approach combining immunophenotypic screens and CITE-seq to link surface immunophenotype and transcriptome. B. Immunophenotypic screens of CB and aBM expression of markers within CD34^+^CD38^-^ with fold change (mean(Abx+% aBM)/mean(Abx+ CB)). C. UMAP of CD34^+^ yBM with 15 clusters. D. Dot plot of gene enrichment within clusters, colors represent mean mRNA expression and size of dot represents fraction of cells with expression. E. ADT gateing strategy for CMP, GMP and MEPs. F. Gated cells from E visualized by contour plots on the CD34^+^ UMAP. G. Molecular clusters within immunophenotypic HSPCs, gated on ADT-levels.

To capture all primitive cell populations within human hematopoiesis, we performed CITE-seq[22] analysis with a panel of 43 surface markers on flow-cytometry sorted CD34+CD38^-^HSCs and on CD34^+^ HSPCs. The CD34^+^ yBM cells were visualized using UMAP (Figure1C) and clustered into 16 groups based on their molecular signals. Furthermore, each cluster was annotated based on expression of cell-type specific marker genes (Figure 1D). For example, MLLT3 have recently been shown to maintain HSC identity during culture [27]. In combination with CRHBP and AVP, MLLT3 was used to annotate HSC populations that could be subdivided into three sub-clusters based on their mapping in relation to downstream progenitors. Moreover, differentiated clusters were defined by well-characterized lineage affiliated genes. MEP-I and -II by increasing levels of GATA1, KLF1 and ITGA2B, Lymphoid primed clusters Ly-I and -II by CCR7 and CLNK, and EBF1, RAG2 and MME, respectively. The granulocyte-macrophage progenitor (GMP) cluster expressed ELANE and MPO, while two clusters with dendritic cell signal, denoted DC-I and DC-II, expressed IRF8 and LY86. Interestingly, several intermediate clusters were detected, annotated as MPPs and numerated I-III based on the fading expression levels of the HSC marker genes. While MPP-I cells mapped adjacent to both the HSC-II and HSC-III clusters and expressed low levels of marker genes of differentiation, the MPP-II cluster neighbored exclusively to HSC-II and displayed simultaneous expression of HSC and MEP marker genes, suggesting a Meg/E lineage bias. In contrast, the MPP-III cluster was distant from the HSC populations in the UMAP and lacked expression of HSC-associated genes, but a gain of gene expression related to myeloid and lymphoid clusters. Thus, this data supports a model of human hematopoiesis where megakaryocyte/erythroid (MegE) specification is an early event, separated from commitment into lympho-myeloid lineages. Finally, two cycling cell clusters (Cyc-I and Cyc-II) were defined based on cell-cycle genes like TOP2A and mKI67.

To compare the CD34^+^ heterogeneity defined by single-cell gene expression in Figure1C with documented immunophenotypic signatures for flow cytometry identification of distinct HSPC populations, antibody derived tags (ADT) read-levels were quantified across the UMAP. Cell surface markers known to specify specific populations cohered with marker gene expression (FigureS1B). Advanced flow cytometry protocols utilize several cell surface markers for prospective isolation of each distinct hematopoietic cell type[1, 3, 5]. To compare immunophenotypically defined cell fractions with mRNA-specified cell clusters, we assessed ADT-derived protein expression using established gating strategies for various HSPC populations (Figure1E-F, FigureS1C-D). While the immunophenotypically defined subpopulations overlapped with their expected gene expression cluster, they all displayed molecular heterogeneity. The lymphoid progenitor populations displayed the highest degree of homogeneity, with the Ly-I cluster (63.9%) or Ly-II cluster (67.0%) dominating the immunophenotypic LMPPs and CLPs respectively. In contrast CMPs, MEPs and MPPs were more heterogeneous. CMPs have previously been suggested to represent a collection of cells with different lineage potentials, rather than one multipotent progenitor population[28, 29]. In our data, 63.2% of immunophenotypically-defined CMPs were composed of cells from the MPP-I-III clusters, which represents populations with different lineage differentiation signatures. Thus, our data supports a model where immunophenotypically defined human CMP population consists of a mixture of cell-types with various lineage potential.

Other established progenitor populations such as CD49f^+^HSCs, CD90^+^HSCs (CD38^-/low^CD90^+^CD45RA^-^), GMP, and MEP also stretched across several clusters. However, this heterogeneity was mainly represented by the continuum of molecularly similar populations at different stages of differentiation, which likely accounts for the functional heterogeneity seen within these populations[3, 5]. Interestingly, CD49f^+^HSCs and CD90^+^HSCs are composed of similar molecular clusters with only a minor difference (CD90^+^HSCs:30.3% vs CD49f^+^HSCs:38.7%) of the most primitive cluster (HSC-I).

### Ageing alters the heterogeneity and molecular programs of human MPPs

The functional effects on the hematopoietic system as a result of ageing includes both changes in lineage output[6, 30] and altered frequency and function of immunophenotypically defined progenitor populations, such as HSCs[4, 6]. To elucidate if these alterations are reflected in cellular heterogeneity or gene expression signatures, additional CITE-seq experiments were performed on CD34^+^ and CD34^+^CD38^-^ populations from CB and aBM (>55 years).

Following CITE-seq analysis, CB and aBM were directly compared to yBM using the mapping feature in Single-Cell dwARFed (Scarf) to define each cell type (Dhapola et al, manuscript in preparation). Projection of CD34+ CB cells showed that all yBM subpopulations were present at birth (Figure2A), however the relative proportions of clusters differed. In CB, the MPP clusters, particularly MPP-I (179.5% of yBM), were overrepresented, while HSC-I (35.1% of yBM) and HSC-II (38.5% of yBM) and the more differentiated progenitor populations; MEP-II (38.3% of yBM), GMP (39.1% of yBM), LY-II (28.7% of yBM) and DC-I (16.3% of yBM) were less frequent (Figure 2B). Interestingly, the differences in HSPC heterogeneity observed in yBM as compared to CB were exaggerated in aBM where the multipotent populations; MPP-I (87.3% of yBM), MPP-II (57.4% of yBM) and MPP-III (69.9% of yBM) were further proportionally reduced and HSC-I (102.3% of yBM), HSC-II (154.9% of yBM), as well as MEP-II (209.0% of yBM), and GMPs (108.6% of yBM) were increased in frequency at the expense of Ly-I (19.0% of yBM) and Ly-II (20.9% of yBM) (Figure2A-B).

**Figure 2.**
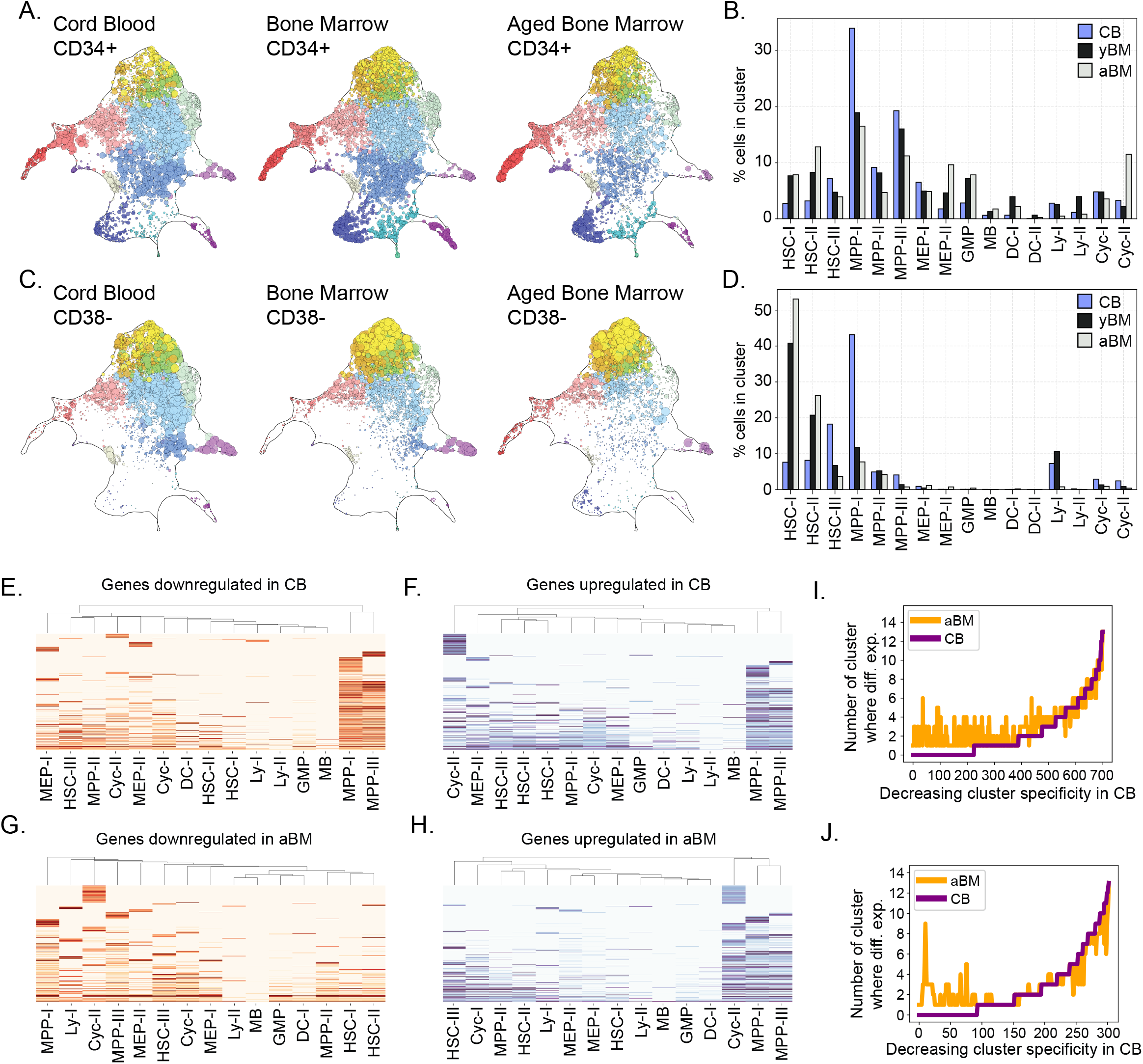
Changes in heterogeneity and gene expression of HSCPCs during ageing. A. Heterogeneity of HSPCs in CB, yBM and aBM projected on CD34^+^ yBM UMAP. B. Quantification of heterogeneity in HSPCs from CB, yBM and aBM. C. Heterogeneity of CD34+CD38-population in CB, yBM and aBM projected on CD34+ yBM UMAP. D. Quantification of heterogeneity in CD34+CD38- for CB, yBM and aBM. E-H. Up- or downregulated genes compared to yBM within scRNA-seq defined clusters. I. Genes (n=669) downregulated in CB and upregulated in aBM vs. yBM. J. Genes (n=288) upregulated in CB and downregulated in aBM vs. yBM.

To improve the resolution of the most primitive populations, CITE-seq analysis was performed on the CD38-cells from each age group (Figure2C-D). Interestingly, while ageing was clearly accompanied with an increased proportion of the most primitive clusters, this effect was unproportional between the HSC-clusters. While HSC-I and –II were relatively expanded 7.1-, and 3.2-fold in aBM compared to CB, the HSC-III cluster was instead 5.1-fold decreased. Meanwhile, the MPP-I was 5.6-fold reduced in aBM and the earliest lymphoid cluster was depleted. Together, this data supports a model where age-dependent hematopoietic changes are manifested in human HSPC heterogeneity, with lineage-balanced-, MPP-dominated hematopoiesis at birth, and MegE-/Myeloid-biased HSC-driven differentiation in the elderly. Moreover, this effect is likely a result of an intra-HSC heterogeneity alteration associated with ageing where HSCs are more prone to differentiate to Meg/E and myeloid lineages (HSC-I and HSC-II) and less to lymphopoiesis (HSC-III).

We further investigated the contribution of the molecularly defined MPPs (MPP-I, -II, -III) within CD123-ADT gated MEP, GMP and CMP populations of CB, yBM and aBM (FigureS1F). Interestingly, within this immunophenotypic CMPs 36.6% and 30.8% consisted of MPP-I cells in CB and yBM, respectively, while MPP-I were 3-fold decreased in aBM. Additionally, within the MEPs, a successive loss of MPP-I was observed, with CB, yBM and aBM being composed of 22.5%, 12.4% and 2.9%, MPP-I cells, respectively. Moreover, when the distribution of MPP-II was investigated, only slight differences between CB and yBM was observed, while aBM MEPs were reduced of MPP-II cells. Finally, within the GMPs, a successive loss of MPP-III cells was observed from CB (42.6%) to yBM (35.5%) and aBM (21.3%). Similar results were also captured when using CD135-ADT to define CMPs, GMPs and MEPs. Together, these results showed that FACS-based protocols for prospective isolation of HSPCs captures populations with different cell compositions, often containing less differentiated cells within CB than yBM and aBM cells. This could at least partly explain the previously observed age-related differences in multipotency[5].

To investigate the molecular programs that correlate with the age-related alterations in cellular heterogeneity, differential gene expression analysis was performed using yBM as reference (Figure 2E-H). Interestingly, while HSC-I and HSC-II populations are transcriptionally stable throughout life, most of the age-related changes in gene expression occur in the lymphoid-related progenitor clusters that are lost over time, such as HSC-III, MPP-I, and MPP-III.

To define whether the observed aged-associated alterations in gene expression in HSPCs were primarily due to intra-cellular transcriptional changes or rather because of changes in cellular composition, genes were ordered according to their cluster specificity. Genes that were downregulated in CB vs yBM, or upregulated in aBM vs yBM (Figure2E), or opposite (Figure2H), were subsequently compared. Interestingly, with decreasing cluster specificity the differentially expressed genes in CB and aBM correlated in both gene expression gained and lost during ageing. Among the genes generally upregulated during ageing were CLU, DNTT, B2M as well as myeloid genes, such as ELANE and MPO, while ERG1, ID2 and KLF2 were downregulated (TableS2). This indicates a gradual ageing phenotype across the cells of the HSPC compartment starting from birth and continuing throughout life. Together, these results suggest that the gene expression alterations of the HSPC population during ageing do not depend on drastic changes in transcriptional programs of individual cells, but rather on changes in cellular composition.

### Identification of an HSC-specific enhancer signature

CITE-seq analysis is powerful in describing heterogeneity based on single-cell gene expression signatures and has the advantage of linking subpopulations to cell-surface proteins. However, ScRNA-seq is inefficient in capturing low-abundant mRNAs like transcription factors (TFs)[19]. In contrast, ScATAC-seq that measures open chromatin, allows for definition of cell-type specific gene regulation by simultaneous prediction of enhancer accessibility, TF-activity, and transcription. Thus, ScATAC-seq provide a higher dimensional and earlier indication of gene activity, which potentially might be useful for detecting the molecular changes underlying lineage priming or bias of HSPCs.

ScATAC-seq was performed on HSPCs, visualized by UMAP (Figure3A) and fractioned into 14 clusters. The clusters were subsequently annotated by assessing the accessibility of lineage specific TF-motifs (FigureS2A). This revealed a similar UMAP structure of the ScATAC-seq data as observed for the ScRNA-seq, including an HSC cluster at the apex of a continuum of progenitor cells that preceded more distinct clusters of myeloid, lymphoid, dendritic, basophilic and erythroid lineages, respectively.

**Figure 3.**
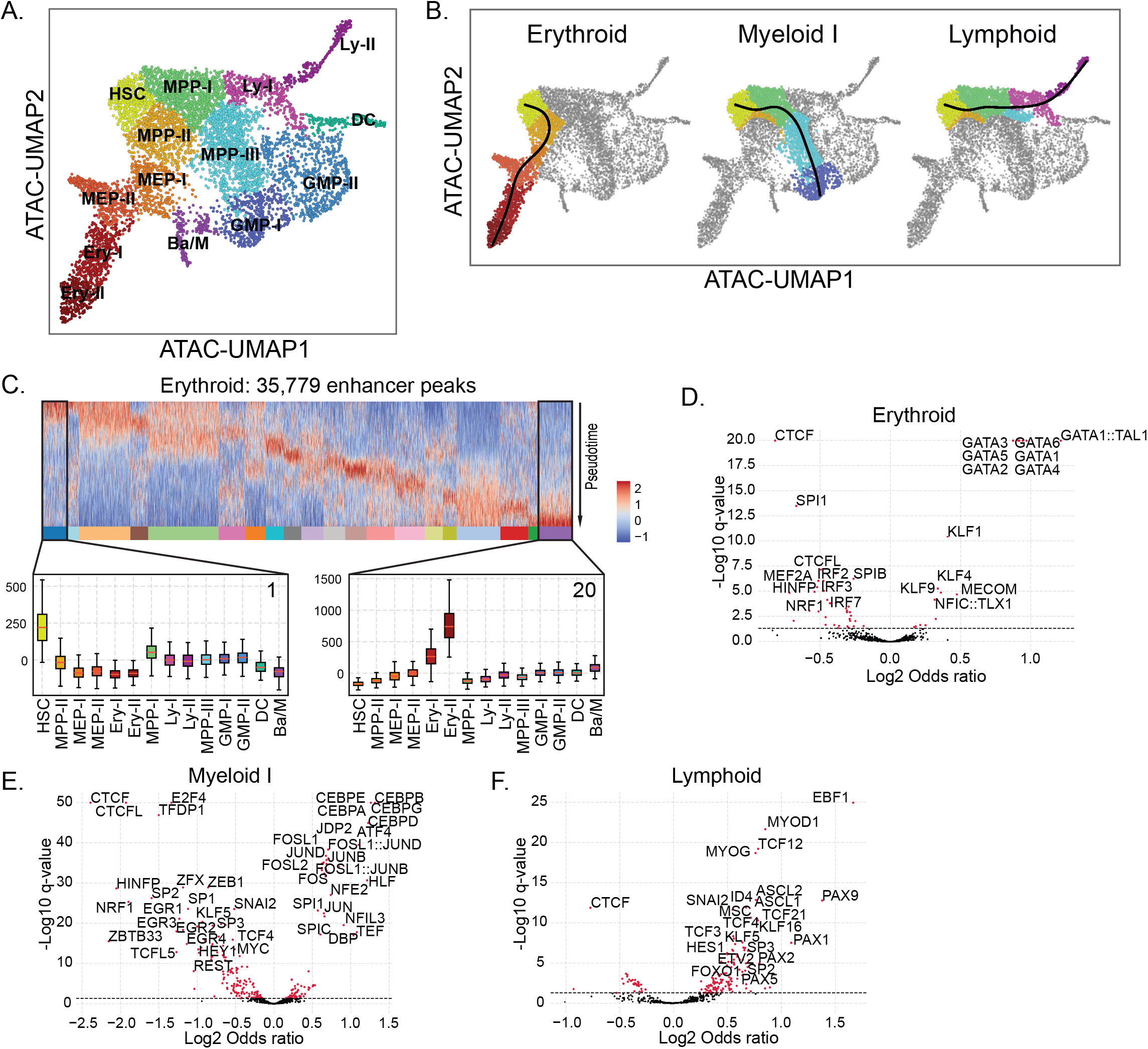
Lineage trajectory analysis of sc-ATAC-seq defines TF-motifs and enhancer gene targets enriched in lineage specific enhancers. A. ScATAC-seq of yBM HSPCs with 14 clusters. B. Lymphoid, Myeloid I and Erythroid differentiation trajectories in yBM CD34+ cells. C. Heatmap of enhancer accessibility along Erythroid lineage trajectory. Enhancer clusters with primitive (enhancer cluster 1) and differentiated (enhancer cluster 20) according to pseudotime are denoted, with boxplots displaying cluster-wise enrichment of enhancer accessibility. D-F. Enhancer enriched TFBS motifs within Erythroid, Myeloid I and Lymphoid terminal ends.

Due to the continuous nature of HSPC differentiation[15, 16], the lineage relationships between cells in the ScATAC-seq was investigated using trajectory analysis where six terminal endpoints were defined. Interestingly, while four lympho/myeloid trajectories followed a similar path through the MPP-I cluster, the Meg/E and Ba/M trajectory diverged early from HSCs into the MPP-II cluster (Figure3B, FigureS2B). Thus, both ScRNAseq and ScATACseq analysis supports a model for human hematopoiesis where Meg/E and lympho/myeloid commitment occurs early in stem cell differentiation.

It is becoming increasingly clear that enhancer activity, compared to promoter accessibility or gene transcription, is a better indicator of cell type specification[20, 21]. To investigate the dynamic changes in enhancer activity along the trajectories within the ScATAC-seq data, we utilized the GeneHancer database[31] to define enhancers and their putative gene targets. In total, 35,801 enhancers were identified in the HSPCs. For each of the six trajectories, the enhancers were ordered along the pesudotime axes (Figure3C, FigureS2B). Next, we set out to define TF activity as measured by TF binding site (TFBS) over-representation within the data. Unsurprisingly, lineage specific TFBS were identified within enhancers at the terminal ends of each trajectory. The lymphoid end was characterized by PAX, TCF and EBF1, the myeloid by CEBPs, JUN, FOS and SPI1, while the Meg/E displayed accessible TFBS for GATA, KLF and TAL1 (Figure3D-F). Intriguingly, the TF CTCF and its paralogue CTCFL, were significantly underrepresented within all terminal ends, suggesting that CTCF activity is incompatible with lineage commitment. Indeed, CTCF has recently been defined as an important regulator of stemness[32].

To integrate the ScATAC-seq data with the ScRNA-seq analysis of yBM HSPCs, we next compared the mRNA expression levels of terminally active TFs of each trajectory (FigureS2D,F,H). Furthermore, the ScRNA-seq data was investigated for mRNA expression of putative gene targets for terminal end enhancers (FigureS2E,G,I). Encouragingly, a strong overlap of both approaches was observed. Thus, integration of ScATAC-seq and ScRNA-seq data allows for sensitive and comprehensive identification of cluster-specific enhancers, relevant TF activity, and enhancer target-gene expression.

Even though the integration of ScRNA- and ScATAc-seq data is efficient in defining transcriptional signatures responsible for lineage identity, the capacity to perform multiomics at single-cell resolution has even more merit for defining molecular states of less distinct populations, such as early progenitor stages and HSCs[33, 34]. Thus, when focusing on early pseudotime, 14,731 early enhancers were defined, with 650 enhancers in common for all six trajectories. These sets of enhancers were denoted as HSC enhancers and HSC-specific enhancers, respectively. The cumulative accessibility of these HSC-specific enhancers was enriched within the most primitive cells (Figure4A-B).

**Figure 4.**
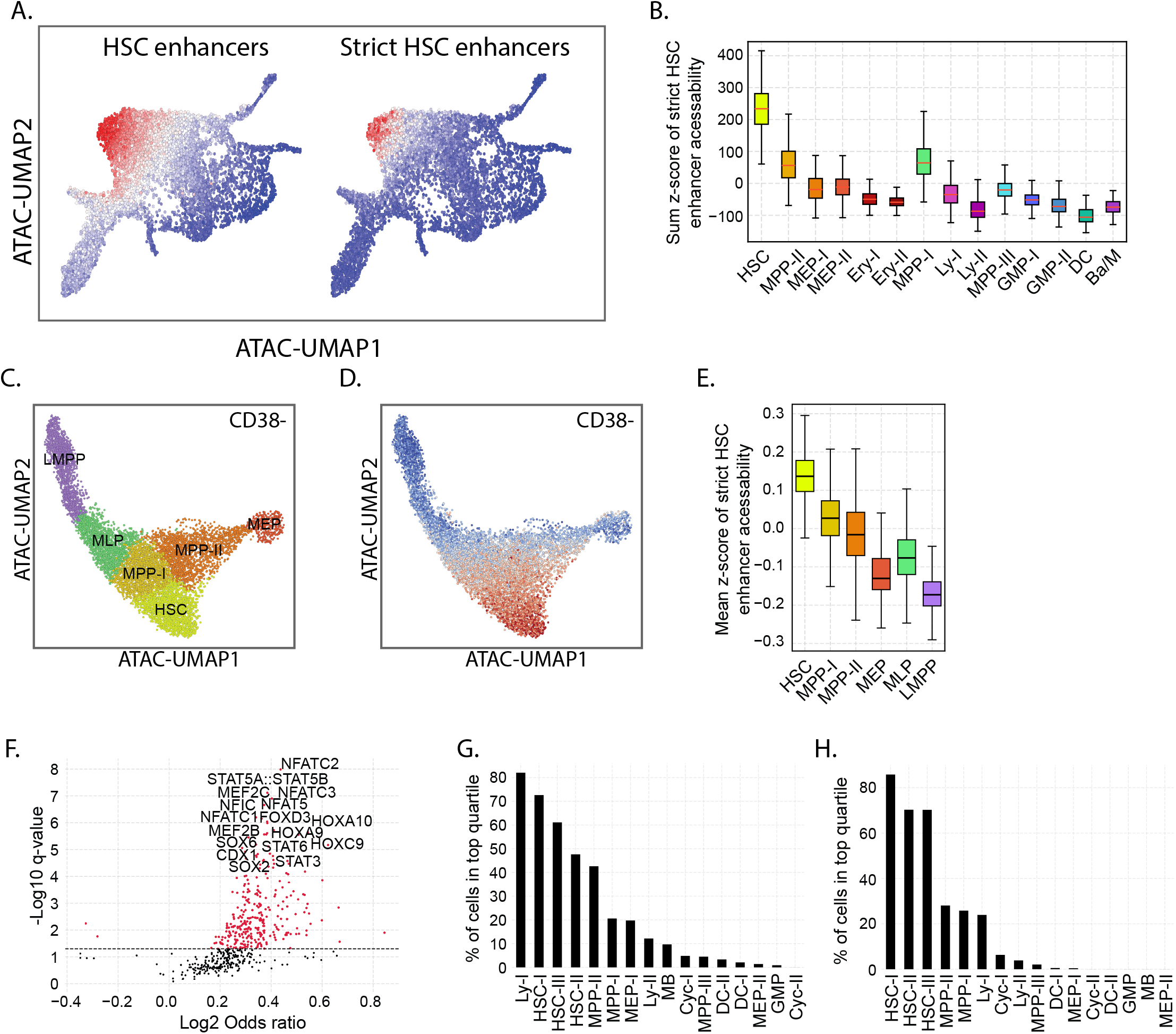
Hematopoietic stem cell specific enhancers defined by lineage trajectory analysis. A. HSC enhancer and strict HSC enhancer accessibility in HSPCs. B. Strict HSC enhancer accessibility within ScATAC-seq clusters. C. ScATAC-seq of Lin-CD34+CD38- sorted cells. D. Accessibility of strict HSC enhancers within CD34+CD38- cells. E. Accessibility of strict HSC enhancers within CD34+CD38-clusters. F. TFBS motifs enriched in strict HSC enhancers. G. Proportion of cells within ScRNA-seq defined clusters with strict HSC enhancer associated TF mRNA values in the top quantile. H. Proportion of cells within ScRNA-seq defined clusters with enhancer target gene mRNA values in the top quantile.

To improve the resolution of the most primitive populations, ScATAC-seq was performed on yBM CD34^+^CD38^-^HSCs cells (FigureS3) and visualized in a separate UMAP divided into six clusters. The clusters were defined as; HSCs, MPPs (MPP-I and MPP-II), myeloid-lymphoid-like progenitors (MLP), LMPPs, and MEPs, based on mapping to CD34^+^ clusters (Figure4C, FigureS3B). Of note the CD34^+^ defined HSC cluster were divided across HSC, MPP-I and MPP-II clusters of the CD34^+^CD38^-^HSCs population, with 50% mapping to the HSCs. This suggests that further separation of the HSC cluster is a result of the higher-resolution obtained within the CD34^+^CD38^-^HSCs- (FigureS3B-C). Next, the accessibility of the HSC-specific enhancer signature was assessed within the CD34^+^CD38^-^HSC population (Figure 4D-E). As expected, the enhancer signature was enriched within the HSC cluster, with a distinct and even loss of accessibility that correlate with differentiation into the two lineage-biased MPP-I and MPP-II subsets.

The identification of a high-resolution, HSC-specific enhancer landscape allows for identification of active TFs as described above. The HSC-specific enhancers revealed a human HSC TF-activity signature dominated by several previously reported TFs important for HSC identity[35-39], such as HOX- (HOXA9, HOXC9 and HOXD13), SOX- (SOX2, SOX3 and SOX6) and STAT family members (STAT1, STAT3 and STAT4) (Figure4F). Additionally, TFs previously not implied to be important for HSC function were observed, including NFAT family members (NFATC1, NFATC2, NFATC3 and NFAT5) (Figure4F). NFAT signaling has been shown to be important for early myeloid differentiation[40] while intracellular calcium levels, an upstream NFAT regulator, have been shown to effect HSC maintenance in-vitro[41].

Similar to the integrated analysis performed on the lineage-specific TFBS, mRNA levels for TFs enriched within the HSC-specific enhancer signature were subsequently measured in the yBM CD34^+^ ScRNAseq. Accordingly, mRNA for TFs with HSC-specific enhancer binding were expressed at relatively higher levels within the HSC-I cluster and decreased in the downstream progeny (Figure4G). Interestingly, upon ranking the clusters from the ScRNA-seq based on frequency of cells expressing the HSC-specific enhancer-binding TFs the Ly-I cluster of cells (LMPPs) expressed the TF-gene signature in comparable frequencies to HSC-I, HSC-II and HSC-III (Figure4G). Indeed, transcriptional similarities between HSCs and LMPPs have previously been documented[42]. However, these results suggest that even though similar TFs are expressed in LMPPs and HSCs, differential enhancer usage results in the functionally distinct phenotypes observed between these two populations. To further explore this hypothesis, the putative target genes of the HSC-specific enhancers was identified, these included HOXC6, KLF6, ITGA6, SOX6, CEBPB and PRDM15 (TableS3). As with the TFs, mRNA expression levels for this HSC enhancer target signature were subsequently measured within the ScRNA-seq (Figure 4H), In clear contrast to TF expression, less than 22% of the cells in cluster Ly-I, where in the top quantile of expression, while the corresponding value for the HSC clusters was approximately 85%. Together this data suggests that HSCs and early lymphoid cells utilize overlapping TFs for their transcriptional control, but these TFs are active in different enhancers, leading to the transcriptional and functional differences observed between the cell types.

### CD35 and CD11A captures the molecular heterogeneity within conventional HSC populations, including bona fide stem cells containing HSC-specific enhancer signature

Prompted by the observation of strong correlations between enhancer usage, enhancer target-gene expression and cell function within the HSPCs interrogated, we next aimed to make use of the CITE-seq data to find a cell-surface marker that allows for capture of the HSC-specific enhancer signature. Pseudotime analysis were applied to the scRNA-seq (Figure5A, FigureS4A) and expression levels of the surface markers were ordered along the pseudotime for each lineage (Figure5B, FigureS4B). Consistent within all differentiation trajectories, cell surface expression of the established HSC marker CD90 was high within the early stages of the pseudotime and faded as differentiation progressed. Additionally, several other cell surface molecules, including CD4, CD54 and CD35, exhibited even stronger expression-bias towards the early stages of the differentiation trajectories. Intriguingly, among all 43 surface markers, CD35 was the only marker with enriched expression (1.2-fold) in the HSC-I cluster as compared to MPP-I cells (FigureS3C). Moreover, visualizing the CD35 expression on the UMAP (Figure5C) confirmed high and specific expression within the HSC clusters across the different ages (CB, yBM and aBM).

**Figure 5.**
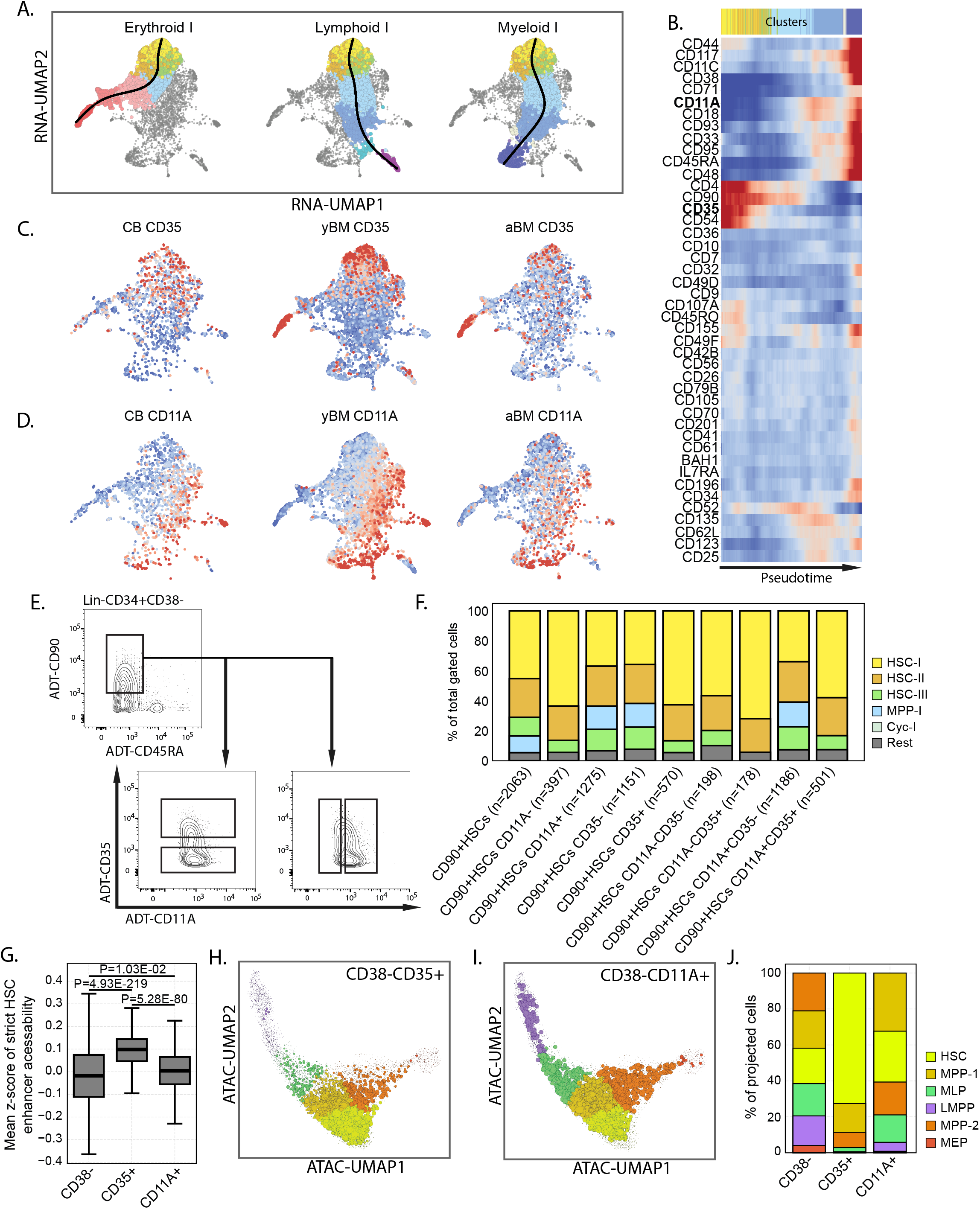
CD35 and CD11A can be used to prospectively isolate HSC Enhancers signature. A. Psuedotime ordering of CD34^+^ yBM CITE-seq, of Erythroid, Myeloid I and Lymphoid I trajectories. B. CITE-seq ADT expression ordered along psuedotime of the Myeloid I trajectory. C-D. CD11A and CD35 protein expression displayed on CB, yBM and aBM UMAP, red corresponds to high values and blue low. E. ADT-gating of CD90^+^HSC, using CD11A and CD35 within sorted CD38^-^HSCs. F. Proportion of ScRNA-seq clusters captured within the gated cells. G. Accessibility of strict HSC enhancers within sorted CD38^-^, CD38^-^CD35^+^ and CD38^-^CD11A^+^ cells. H-I. Projection of CD38^-^CD35^+^ and CD38^-^CD11A^+^ cells on CD38^-^ ScATAC-seq UMAP. J. Proportion of cells within clusters from CD38^-^, CD38^-^CD35^+^ and CD38^-^CD11A^+^ cells.

In contrast, cell-surface expression of CD11A, previously reported as a negative marker for murine HSCs[24], displayed an opposite pattern compared to CD35, with low abundance within the HSC-I cluster and increasing expression along the pseudotime within the lymphoid or myeloid differentiation trajectories (Figure5B, FigureS4B). Furthermore, visualization of CD11A expression in the ScRNA-seq UMAP suggested that CD11A could be used as a negative marker of human HSCs and mark early lympho-myeloid lineage priming at all stages of life (Figure 5D).

To explore the potential for CD35 and CD11a to enrich for stem cell molecular signatures within HSC containing populations, advanced multiparameter gating was performed using ADT expression from the yBM CD34^+^CD38^-^HSC CITE-seq data. Current state-of-the-art protocols for HSC isolation, CD49f^+^HSCs or CD90^+^HSCs, was further divided by different combinations of CD35 and CD11a and the resulting cell fraction was examined for relative proportion of ScRNAseq-defined clusters (Figure5E).

Surprisingly, despite the well-established enrichment in reconstitution activity obtained within CD49f^+^HSCs, the CD49f^+^HSC and CD90^+^HSC populations constituted highly similar heterogeneity of cells from the clusters with the most primitive molecular signatures (e.g. 45.1% and 45.2% HSC-I, 27.2% and 25.8% HSC-II, respectively) (Figure5F, FigureS4D). Instead, dividing the CD90^+^HSC populations using CD35 and CD11A had substantial effect on the cluster composition. The CD35^+^ cells were depleted from MPPs, entirely consisting of cells from HSC-I (62.6%), HSC-II (24.0%), and HSC-III (7.9%) (Figure5F). Gating of the CD11A^-^ population had a similar effect as CD35 by enriching for cells with HSC-like transcriptional signatures. Interestingly, the combination of both markers allowed for isolation of a CD35^+^CD11A^-^ population. This was 1.6-fold enriched for the HSC-I cluster compared to the CD90^+^HSCs, and depleted of HSC-III cells, thus exclusively consisting of HSC-I (71.9%) and HSC-II (22.5%) cells (Figure5F). These results were recaptured by using CD49f^+^HSCs together with CD35 and CD11A, which again showed enrichment of HSC-I (FigureS4D). In conclusion, gating of CD35^+^ and CD11A^-^ cells within conventional state-of-the-art HSC populations purifies the most primitive molecular signatures observed in HSPC ScRNA-seq data.

Next, we explored the capacity of CD35 and CD11A to capture the HSC-specific enhancer signature. Lin^-^CD34^+^CD38^-^CD35^+^ and Lin-CD34^+^CD38^-^CD11A^+^ yBM cells were sorted and subjected to scATAC-seq. The accessibility of the strict HSC enhancer signature was assessed in each sorted population. Despite the effects observed from the combination of both markers on mRNA-defined heterogeneity, only CD35^+^ sorted cells showed a significant enrichment of HSC-specific enhancer activity when compared to the CD38^-^ population (P-value 4.93E-219, students t-test, Figure5G), while the HSC-specific enhancer signature was unchanged in CD11A^+^ cells (P-value 1.03E-2, students t-test).

To investigate which ScATAC-seq-defined subpopulations are enriched within the CD35^+^ and CD11A^+^ cells, the populations were projected on the CD34^+^CD38^-^ UMAP (Figure5H-J). CD11A^+^ cells were enriched in both the MPP-I and HSC populations, with 30% of the cells mapping to the MPP-I and 30% mapping to HSC clusters, representing a 1.5-fold enrichment over the CD38^-^ cells. However, there was a distinct loss of MEP clusters within the CD11A^+^ population, supporting the CD11A lympho-myeloid bias observed in the CITE-seq analysis. Moreover, CD35^+^ cells showed a marked increase of the HSC population, with 70% of the cells mapping to the HSC cluster, which represented a 3.5-fold enrichment compared to the CD34^+^CD38^-^ population. The increased capture of HSCs by CD35 was at the expense of differentiated populations, with MPP-I and MPP-II constituting only 20.5% and 10% of the CD35+ population, respectively. Meanwhile, the other clusters (MEP, MLP and LMPP) were depleted, further validating CD35 as a marker of epigenetically primitive cells.

To explore the relevance for the HSC-specific enhancer signature, CD35 and CD11A were exploited in FACS protocols to subdivide the HSC population for functional studies. First, FACS analysis of CD35 and CD11A on CB cells together with conventional markers revealed a distinct and consistent CD11A/CD35 profile that subdivided the CD34^+^CD38^-^HSCs, CD90^+^HSCs and CD49f^+^HSCs, populations in all donors(Figure6A). Enrichment of CD35^+^ cells was observed with each sequential step of putative HSC purification, culminating with HSC-specific enhancer signature-containing CD35^+^ cells making up 60%, and CD35^+^CD11A^-^ cells 15% of the CD49f^+^HSCs (Figure6B).

**Figure 6.**
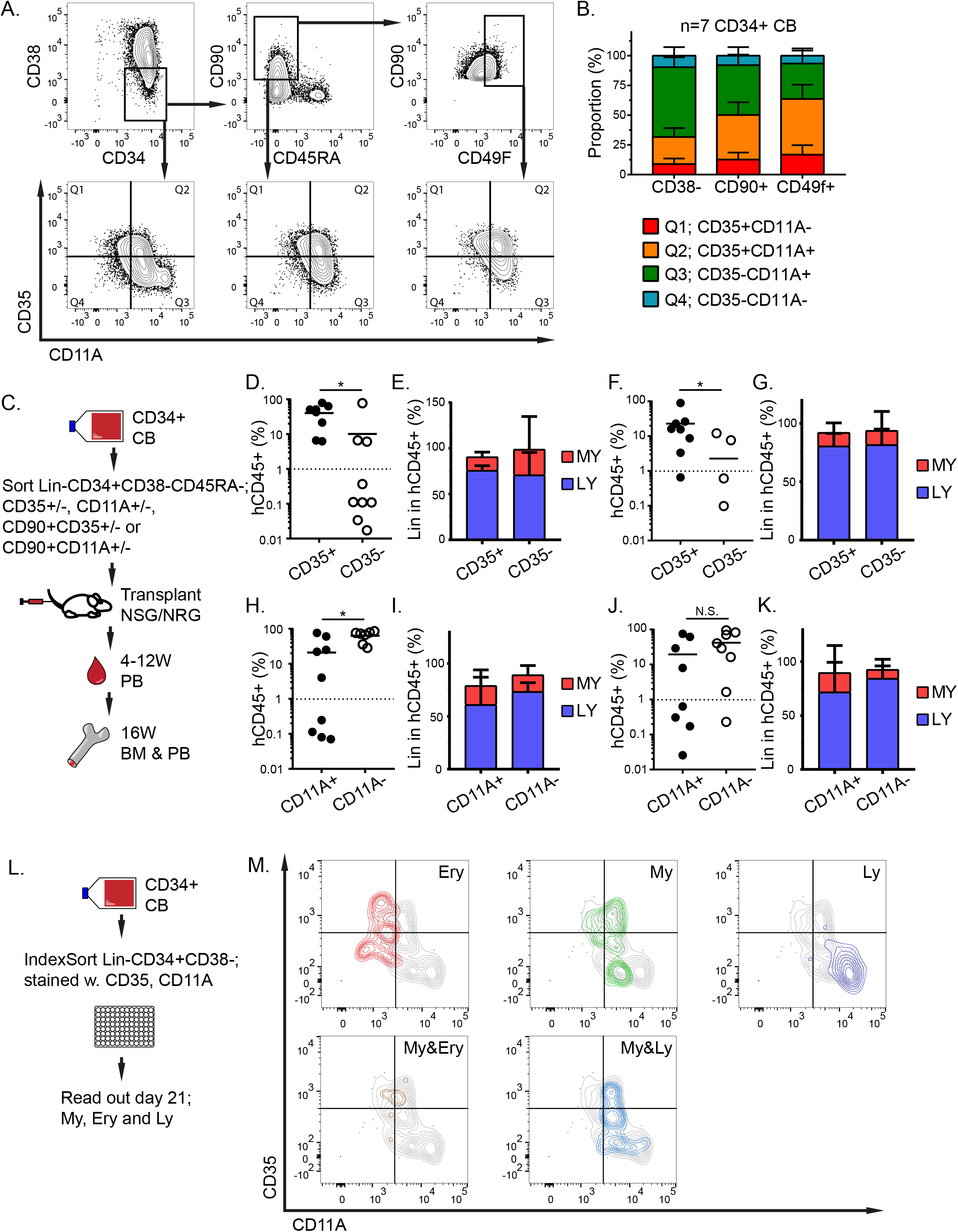
CD35 and CD11A confirms function of HSC enhancer signature. A. Representative FACS plots of CD35 and CD11A expression within primitive immunophenotypic populations in CB. B. Summary of fraction of CD35 and CD11A in seven CB-donors. C. Experimental layout of transplantation experiments. D-E. % human CD45 engraftment and lineage reconstitution of 1000 Lin^-^CD34^+^CD38^-^CD45RA^-^CD35^+/-^ cells in BM after 16W. F-G. % human CD45 engraftment and lineage reconstitution of 500 Lin^-^CD34^+^CD38^-^CD45RA^-^CD90^+^CD35^+/-^ cells in BM after 16W. H. % human CD45 engraftment and lineage reconstitution of 1000 Lin^-^CD34^+^CD38^-^CD45RA^-^CD11A^+/-^ cells in BM after 16W. J-K. % human CD45 engraftment and lineage reconstitution of 500 Lin^-^CD34^+^CD38^-^ CD45RA^-^CD90^+^CD11A^+/-^ cells in BM after 16WL. Experimental layout of in-vitro single cell assay. M. Index-sorted lineage potential of Lin^-^CD34^+^CD38^-^ single cells differentiating to erythroid (Ery), myeloid (My), lymphoid (Ly), myelo-erythroid (Ery&My) and lympho-myeloid (My&Ly).

Next, Lin^-^CD34^+^CD38^-^CD45RA^-^ HSC-enriched cells or CD90^+^HSCs were subdivided by CD35 or CD11A and all resulting cell fractions were separately xenotransplanted into NSG or NRG mice (Figure6C). Intriguingly, CD35-enrichment for cells characterized by the HSC-specific enhancer signature clearly discriminated between multilineage repopulation activity at 16 weeks post transplantation. In both Lin^-^CD34^+^CD38^-^CD45RA^-^ (Figure6D-E) and CD90^+^HSC populations (Figure6F-G), prospective isolation of CD35^+^ cells resulted in a 4.0-fold and 10-fold increase in human engraftment compared to CD35^-^ cells, respectively. While 15 out 16 recipients transplanted with CD35^+^ HSCs were repopulated (>1% hCD45^+^) with human cells, only 5 out of 18 mice receiving CD35^-^ donor cells displayed human engraftment (>1% hCD45^+^). Importantly, the human CD34+ HSPC compartment was exclusively reconstituted by CD35^+^ HSCs (FigureS5C). Thus, the HSC-specific enhancer signature captured by CD35 is critical for human multilineage reconstitution potential and self-renewal in xenotransplantation models.

In agreement with the less distinct enhancer signature observed when using CD11A, both the CD11A^-^ and the CD11A^+^ HSCs contained SCID repopulating activity (Figure6H, J). However, reconstitution capacity of both mature cells and CD34^+^ HSPCs were significantly higher for CD11A^-^ HSCs compared to CD11A^+^ cells.

Since human xenograft models have a predisposition for lymphoid output[43], single cell in-vitro cultures were performed to further investigate the lineage capacity of the CD35 and CD11A HSC populations. CB CD38^-^HSCs were index-sorted for CD11A and CD35 expression into wells supporting erythroid, myeloid and lymphoid (NK- and B cell) differentiation. Following 21 days, the output from each clone was analyzed (FigureS5D) and correlated to indexed FACS data for CD11A and CD35 (Figure6M). While CD35/CD11A expression had little effect on the cloning efficiency (FigureS5E), erythroid differentiation potential was significantly enriched for the CD11A^-/low^ population, while CD11A^+^ cells were increasingly biased for lympho-myeloid differentiation (Figure 6M, FigureS5F). This is in accordance with the CITE-seq data, were the CD11A expression correlated with lympho-myeloid pseudotime, and supports the trajectory analysis that indicated a split of MegE and Lympho-myeloid commitment already at the stage of HSC-II and HSC-III clusters of the Sc-RNAseq (Figure5A, D). Notably, all these clusters are captured within CLOUD-HSPCs and current state-of-the-art FACS protocols for prospective isolation of human HSCs (FigureS1C-D), demonstrating the functionally relevant heterogeneity within immunophenotypic HSC populations. In contrast, CD35^+^ cells differentiated into all lineages, including potent erythroid output (Figure 6M, FigureS5G). Again, this is in line with the identification of an HSC-specific enhancer signature within CD35^+^ HSCs, as well as supportive of MegE potential being specified within the most primitive subpopulation of HSC fractions.

Taken together, our functional experiments validated observations from CITE-seq and ScATAC-seq analysis by demonstrating that bona fide HSCs are characterized by an HSC-specific enhancer signature critical for multilineage differentiation capacity, SCID-repopulating activity, and self-renewal. This subtype of HSCs can be captured from conventional HSC-containing populations based on CD35 and CD11A expression.

## Discussion

Traditionally, the heterogeneity of hematopoietic progenitor cells has been approached by retrospective definition of cell-type composition following functional read-outs of prospectively isolated cell fractions. This design, including distinct FACS gating, measurement of functional capacity rather than fate, and generalization of functional output from preconceived populations, inherently favors the step-wise hierarchical model for hematopoietic differentiation. Recently, re-analysis of the HSPC compartment by ScRNA-seq suggested that the CD34^+^CD38^-^ progenitors of hematopoietic differentiation are unclusterable, forming a continuum of low-primed undifferentiated hematopoietic stem and progenitor cells that gradually acquire transcriptomic lineage priming, resulting in the production of unilineage-restricted cells and loss of self-renewal capacity[15, 16]. Thus, according to the CLOUD-HSPC model, the subsequent progeny of HSCs (e.g. MPPs) would not represent distinct cell types, but rather CLOUD-HSPCs in a transitory state towards a certain lineage. Indeed, our data combining ScRNA-seq and ScATAC-seq analysis support the CLOUD-HSPC model in that the earliest stages of the hematopoietic hierarchy does not subdivide into distinct clusters but rather form one molecularly similar population at the apex of committed progenitors. However, trajectory analysis suggested that a bias towards either Meg/E or Lympho/Myeloid fates occurs within the HSPC population immediately downstream HSCs. This was especially apparent when trajectories were defined based on enhancer accessibility. Importantly, CITE-seq data revealed that cell surface expression of CD11A tightly correlated with lympho-myeloid differentiation and was upregulated already at this early point of trajectory separation within HSPCs. Even though little or no lineage priming could be observed on chromatin or mRNA level at this point, functional experiments revealed that CD11A expression even within highly purified HSPC populations is incompatible with erythroid differentiation. Thus, our data supports a CLOUD-HSPC model where, despite subtle differences in global gene expression, distinct lineage-biased progenitor types exist and can be prospectively isolated.

Since accumulating data indicate that the function of a cell is reflected primarily by its enhancer accessibility[19, 20], we reasoned that a fraction of the CLOUD-HSPCs should contain an HSC enhancer signature that is decisive for HSC function. Our analysis provided an epigenetic signature consisting of 650 enhancers specifically accessible within HSCs and rapidly closed as differentiation trajectories are established. By TFBS analysis, these enhancers were overrepresented for established HSC TF motifs, including HOX, SOX and STATs[35-39]. Interestingly, novel candidates like the NFAT family of TFs were additionally enriched within the HSC enhancers. NFATs does not have a documented role in HSCs, but have been implicated in lympho-myeloid lineage decisions[40], making them an interesting target for further investigations into lineage priming.

Interestingly, gene expression levels for HSC-specific enhancer-enriched TFs were promiscuous and enriched within HSCs, MPP as well as MLP (LMPP) populations. Although the transcriptional similarities of these populations have been shown before[42], our data demonstrate how similarities in TF expression result in differences in cell-specification. By using an enhancer centric approach, we observed that TFs overlapping in both HSCs and MLPs interact with different enhancers, subsequently driving expression of different genes. This observation has implications for analysis of lineage priming prompting for enhancer accessibility as markers of primed states, ahead of conventional definitions based on co-occurrence of lineage specific and multipotency defining TF mRNAs.

Using the ADT-data from CITE-seq we could identify cell surface markers that correlated with HSC-specific enhancer target gene expression. CD35^+^ cells captured the most primitive state of each lineage trajectory according to ScRNA-seq data, and were superior to all conventional markers in enriching for the HSC fraction as defined by ScATAC-seq, especially when used in combination with CD11A that marks lympho-myeloid biased HSPCs. Importantly, CD35^+^ isolation resulted in a multifold and significant purification of the HSC-specific enhancer signature compared to unfractionated CLOUD-HSPCs. Most importantly, all functional stem cell activity was captured by CD35, validating that a distinct HSC-specific enhancer signature is tightly linked to functional HSC identity.

The heterogeneity within the most primitive HSPCs observed here are interesting from an ageing perspective, which is characterized by a lymphoid-bias at birth to a myeloid-bias in the elderly [6-8]. Additionally, the frequency of immunophenotypic HSCs increases with age[6, 7, 9]. As a readily available source of HSPCs, neonatal cord blood has been the paradigm for human hematopoiesis, including immunophenotypic definition of HSCs. Here CITE-seq analysis was performed on three different stages of life (CB, yBM and aBM) and age-related changes in immunophenotype, transcription and heterogeneity were directly compared. Intriguingly, the effect of ageing on hematopoietic heterogeneity was not exclusive to the relative increase of HSC frequency, but also associated with a substantial reduction of MPPs, an increased Meg/E and myeloid output, and a reduced number of lymphoid progenitors. Surprisingly, when comparing the gene signatures between the corresponding clusters at different stages of life, we observed that while most progenitor populations, including the HSCs, only experienced minor age-related changes to their gene expression program, substantial transcriptional changes were observed at the lympho-myeloid biased MPP-I and MPP-III stages. Thus, the age-related shift from lineage-balanced to myeloid-biased differentiation is associated with molecular changes within the MPPs rather than HSCs. This finding is consistent with results suggesting that progenitors lose multipotency as they age[5]. Moreover, by ranking differential gene expression of CB and aBM based on their occurrence in clusters, a continuous acquisition of an aged gene expression profile from CB, through yBM to aBM was observed. This gradual development of ageing mirrors recent findings within the mouse system, where scRNA-seq and epigenetic analysis have shown that the foetal to adult transition is a continuous process[44]. Thus, HSPCs are constantly ageing, resulting in global effects on gene expression and cell population heterogeneities.

Taken together, by combining and integrating multi-omic data, we here define the heterogeneity within HSPCs at different stages of life. Trajectory analysis allowed for prospective isolation of lineage-biased cell states within the CLOUD-HSPCs as well as the identification of an HSC-specific enhancer signature tightly linked to stemness. These results advance our understanding of human haematopoiesis and position enhancers as the gatekeepers of hematopoietic cell identity. Future studies will evaluate the importance of stem cell-related enhancer programs also in disease settings i.e., leukaemia, and their therapeutic potential.

## Methods

### Sample preparation

Human cord blood and bone marrow was obtained from either consenting mothers or donors and processed according to guidelines approved by Lund University. In brief, cord blood (CB) or bone marrow (BM) was diluted 1:1 with PBS (GE lifescience) with the addition of 2% fetal bovine serum (GE lifescience) and 2 mM EDTA (Ambion), added to lymphoprep tubes (Alare Technologies) where density centrifugation was performed for 20 min at 800xg. The mononuclear layer was collected, washed 1:1 with Iscove’s modified Dulbecco’s medium (IMDM, Thermofisher) and cells was either frozen down as mononuclear cells (MNCs) or processed further for CD34 enrichment. CD34 enrichment was preformed using CD34 enrichment kit (Miltenyi Biotec) according to manufacturer’s protocols, in brief, cells were stained with FC block and CD34 enrichment beads for 20 min, after which on column positive magnetic enrichment was performed. After either MNC or CD34 enrichment all cells were frozen in FBS with 10% DMSO and kept at −150°C until use.

### CITE-seq sample preparation

Cellular indexing of transcriptomes and epitopes was performed according to previous publication [22] with minor alterations. In brief, either MNCs or CD34 enriched cells were thawed and washed and stained with FC block (Miltenyi Biotec) for 10 min and washed again. After FC block cells were stained for 30 min with antibodies for Lineage (CD14, CD16, CD19, CD2, CD3 and CD235a), CD34-FITC and the first set of CITE-seq antibodies (TableS1), cells were then washed before a second stain for 30 min with CITE-seq antibodies, sample specific Hashing antibodies (TableS1) and either a CD38-PECy7 FACS antibody or a CD38-CITE-seq antibody. After the second staining the cells were washed again before being taken up in PBS with 2% FBS and 1/100 7AAD. Cells were then sorted with a FACS AriaIIu (BD), approximately 11,000 cells were sorted from either Lin-CD34+ or Lin-CD34+CD38-populations, all cells were pooled, and single-cell sequencing was done using 10x genomics 3’ V2. Approximately 10,000-40,000 cells were loaded on each channel of the 10x after which the single-cell libraries made according to manufacturer’s instructions, except for the CITE-seq protocol addition. Were instead a primer is added in the cDNA amplification step and instead of discarding the small fragments in the bead purification they are kept for CITE-seq and HASH-seq protocols. The CITE-seq and HASH-seq libraries was prepared according to the publication[22], briefly, the small fragments from the cDNA amplification are purified and the library is barcoded and amplified using PCR with the KAPA high fidelity PCR kit(KAPA Biosystems). After all libraries are ready their quality and size are measured using the bioanalyzer before being sequenced on a NEXTseq (Illumina).

### scATAC-seq

Single cell Assay for Transposase-Accessible Chromatin using sequencing (sc-ATAC-seq) was performed using the 10x genomics single cell ATAC platform according to manufacturer’s instructions. In brief, yBM CD34+ cells were thawed and stained for either, Lin, CD38, CD34 or Lin, CD38, CD34, CD90, CD45RA, CD35 and CD11A. After staining 10,000-40,000 cells were sorted from yBM with four different populations; Lin-CD34+, Lin-CD34+CD38-, Lin-CD34+CD38-CD35+ or Lin-CD34+CD38-CD11A+. After sorting nuclei was isolated by following the 10x low input nuclei isolation protocol, briefly, cells were lysed in lysis buffer (Tris-HCl (pH 7.4, 10mM), NaCl (10mM), MgCl2 (3mM), Tween-20 (0.1%), Nonidet P40 substitute (0.1%), Digitonin (0.01%), BSA (1%), Nuclease free water) for 3 min, washed using washing buffer (Tris-HCl (pH 7.4, 10mM), NaCl (10mM), MgCl2 (3mM), BSA (1%), Tween-20 (0.1%), Nuclease free water), centrifuged for 5 min at 500xg and taken up in diluted nuclei isolation buffer (10x genomics) and centrifuged again for 5 min at 500xg before being taken up in 7µl of diluted nuclei buffer. Nuclei were counted and appropriate nuclei numbers were added to the Tn5 digestion reaction. After digestion nuclei were loaded on to the 10x genomics platform and sc-ATAC-seq were performed. After library preparation the libraries were sequenced on a NOVAseq (Illumina).

### Antibody screens, FACS analysis and sorting

Antibody screens was performed on approximately 300 million CB MNCs, cells were stained with a lineage cocktail (CD14, CD16, CD19, CD56, CD2, CD3, CD123 and CD235a), CD34-FITC, CD38-APC, CD45RA-PB, CD90-BV605 and CD49F-PECY7. Cells were then distributed into LEGENDscreens (Biolegend) in four 96 well plates containing 340 antibodies in separate wells. After staining the screens were run on a FACS canto (BD). Cord blood CD34+ cells were thawed and stained with a lineage-cocktail (CD14, CD16, CD19, CD56, CD2, CD3 and CD235a), CD34-FITC, CD38-PECY7, CD45RA-BV421, CD90-PE, CD35-BV510 and CD11A-BV705. 1000 cells per mouse from 6 populations (Lin-CD34+CD38-CD45RA-CD90+, Lin-CD34+CD38-CD45RA-CD90-, Lin-CD34+CD38-CD45RA-CD35+, Lin-CD34+CD38-CD45RA-CD35-, Lin-CD34+CD38-CD45RA-CD11A+ and Lin-CD34+CD38-CD45RA-CD11A-) and 500 cells per mouse from 4 populations (Lin-CD34+CD38-CD45RA-CD35+, Lin-CD34+CD38-CD45RA-CD35-, Lin-CD34+CD38-CD45RA-CD11A+ and Lin-CD34+CD38-CD45RA-CD11A-) was sorted using a AriaIII/AriaIIu (BD) with a purity >90% into sterile filtered PBS with 2% FBS.

### Transplantations and in-vivo

All animals were processed according to Lund university ethics committee, NSG (Jackson laboratories) or NRG (Jackson laboratories) mice with ages 8-12 weeks was irradiated with 250 cGY and transplanted with the cells, positive controls of 10,000 CD34+ CB cells and untransplanted controls were included.

Peripheral blood (PB) was analysed every four weeks until 16w when BM was analysed. PB was stained with mouse CD45-FITC, human antibodies CD45-APC, CD15-PE, CD33-PE and CD19-BV605. Tibia and femur were collected from both legs and the BM were stained with mouse CD45-AF700, human antibodies CD45-APC, CD15-PE, CD33-PE, CD34-FITC and CD19-BV605, all PB and BM was analysed for human engraftment on an LSR Fortessa (BD). FACS data was then analysed in FlowJo(BD) to determine reconstitution and lineage output, the data was then visualized and analysed in graph pad prism where students t-test was used to determine significance.

### In-vitro assay

For single cell in-vitro lineage read-out, in short, Lin-CD34+CD38-single cells from CD34+ CB stained with Lin, CD34, CD38, CD90, CD45RA, CD11A, CD35, CD49f and CD45RO were index sorted into flat-bottom 96-well plates. Plates pre-coated with 0.1% gelatine, were seeded with 3000 MS5 stromal cells/well in 100µl OptiMEM (ThemoFisher) on the day before sorting and kept in 37°C overnight in a humidified incubator. On the day of sorting the OptiMEM was removed and instead 50µl culture media (Myelocult H5100 (StemCell technologies) supplemented with SCF (100ng/ml), TPO (50ng/ml), FLT3L (50ng/ml), IL-6 (20ng/ml), IL-2 (10ng/ml), IL-7 (20ng/ml), EPO (3u/ml) and PEN/STREP (1%)) was added. After 4 days 50µl of culture media supplemented with 2X cytokines was added and after 14 days from experiment start 100µl of culture media supplemented with 2X cytokines was added. After 21 days all cells were transferred to V-bottom 96-well plates, washed and stained with CD235a-FITC, CD19-PE, CD11b-APC, CD45-AF700, Nkp46-BV421, CD33-BV785, CD14-BUV395 and CD71-BUV496. After staining for 45 min at 4°C cells were washed and taken up in 200 µl PBS+2%FBS and 1/1000 7AAD before being run on a LSR Fortessa (BD). Lineage out-put was analysed in FlowJo, number of cells in gates were then exported into csv file format and loaded into R. In R index sorted cell identities were linked to lineage out-put through Flowcore, to enable linking lineage potential to cell identity. Data was then visualized using FlowJo and graph pad prism.

### Bioinformatic Method

#### CITE-seq analysis

Post sequencing libraries were demultiplexed using Cellranger version 3.0.2, and loaded into Seurat[45] (ver. 3.0.1). Cells that did not have more than 50% of all HTO UMIs from only one HTO were discarded as doublets/multiplets. The remaining cells were assigned an HTO/sample identity based on the HTO index that had more than 50% of UMIs for that cell. yBM HSPCs with UMI counts outside the range 2000 and 35000 were removed. Also, only those cells that had a number of detected genes within 5000 and 4500 were retained. Cells with mitochondrial gene UMIs percentage higher than 5% were removed. Filtering criteria’s for the rest of the samples can be found in the sup. table 4. Post-filtering the yBM CD34+ cells were log normalized, and 2000 highly variable genes were identified individually from each replicate sample. The common HVGs (1206) between the replicates were used to perform the PCA (principal component analysis) reduction and the 15 top PCs which were used to construct an UMAP embedding of the cells. The cells were then clustered using the default clustering algorithm in Seurat with resolution parameter set at 0.88, yielding 13 clusters. The ‘FindAllMarkers’ function in Seurat were used to identify positive marker genes (which expressed in at least 25% of cells in a given cluster) for each of the clusters. Cluster identities were then designated based on these marker genes. The HSC was further subdivided into three subclusters using only the 1063 cells from the HSC cluster. This subset of cells was sub-clustered (resolution=0.3) using Seurat using top 1000 HVGs and four PCA dimensions.

#### Pseudotime analysis CITE-seq

Trajectory analysis was performed using Slingshot (Bioconductor version 3.9), in short, the umap and cluster identities defined using Seurat were loaded into Slingshot. By using the HSC-I cluster as the starting point and the default parameters, Slingshot defined six trajectories traversing the intermediate cell states.

#### ADT Gating

ADT expression was normalized using the CLR (centered-log ratio) normalization in Seurat. Prior to exporting the normalized values into FCS format, the values were transformed by calculating their antilogs (base e) and multiplied by a scaling coefficient (1000). The FCS files were loaded into FlowJO software for gating. Gates were set using internal negative controls to define conventional and novel populations. The cells within each gate were then exported using FlowJO export into CSV files, which were then read into python and the cell identities were defined by matching exported ADT expression to original ADT levels of cells.

#### Projection CB, yBM and aBM

Projection of CD34+/CD38-cell populations of CB, yBM and aBM was done using Scarf (version 0.4.4). In brief, the yBM CD34+ UMAP, HVGs and cluster information from the Seurat analysis was loaded into Scarf, and by using the “run_mapping” function (n neighbours = 9 and PC dimension = 15), the nearest neighbours of projected cells in the yBM population were identified. Based on the closest molecular match the cells are classified into individual clusters using the ‘get_target_classes’ function (threshold=0.4).

The projection mapping scores for a given sample were calculated using ‘get_mapping_score’. The size of the yBM cells was set proportional to the mapping scores in the UMAP to indicate the degree of projection (weighted number of nearest neighbours) onto each cell. The proxy location of projected cells on the UMAP was made by using the coordinates for the nearest neighbours of the projected cell among the yBM cells.

#### Differential gene expression analysis

After cells were classified into clusters using Scarf, each cluster and sample was pseudo-bulked into three replicates. To perform pseudobulking, for each cluster, cells were randomly partitioned into three groups. Then the sum of expression values of each gene was calculated within the randomly constructed groups. Then using DEseq2[46], CB and aBM clusters were compared to their corresponding clusters in yBM population. Genes which were significantly up- or down-regulated (p-value <0.001 and absolute fold change > 2) were used for further analysis.

### scATAC-seq analysis

#### Barcode filtering and MACS2 peak calling

##### Barcode filtering

Post-sequencing nuclei were processed using the cellranger-atac pipeline (version 1.2.0), after alignment to the human genome (GRCh38) and barcode identification. Each barcode identified by Cellranger in the aligned reads are categorized into either cell associated barcodes or background barcodes. To perform this categorization, a sorted list of the total number of UMIs (Log2 transformed) associated with each barcode were created. Thereafter the minima in the gradient of this list were calculated using Numpy’s ‘gradient’ function. The minima in the first 100 and beyond 10000th elements are ignored because the expected number of cells were between that range. Cells were further removed if they had had either too high (log2 % > 4) or too low (log2 % < −3) percentage of reads from mitochondrial genes.

##### MACS2 peak calling

Only the fragments were recovered from the BAM by considering only those reads that aligned as proper pairs and with mapping quality (MAPQ score) higher than 20. Fragments associated with non-cell barcodes were discarded. The terminal ends of the fragments were saved thus recording all the cut sites in the BAM files. The chromosome-wise BED files of cut sites were merged from the replicates and only one cut site was recorded at a given genomic location. The sorted, replicate merged, chromosome-wise BED files were then used to identify regions of high accessibility (‘peaks’) using MACS2 software. For each population of cells the same parameters used in the MACS2 call were: ‘-s 150 --nomodel --shift −75 --extsize 150’; the genome size was set as chromosome size in each MACS2 call. Next, in order to create a cell-peak matrix of each population, the number of cut sites in each peak for each cell were recounted. Additionally, the cut-sites of the samples for the peak regions of other samples were also counted. For example, the cut-sites from yBM CD38-cells were taken and recounted in the yBM CD34+peaks to create a cell-peak matrix of yBM CD38-in CD34+ defined feature set (peaks).

##### Peak filtering

The cell-peak matrices for each population were then processed using Scarf to perform another round of cell and feature filtering. Peaks that existed in the ENCODE-defined blacklisted regions (version 2) and those from X and Y chromosomes were removed.

#### Gene Score and enhancer matrix calculation and cell filtering for yBM CD34+ cells

##### Gene score calculation

The gene scores for each cell were thereafter calculated. The score for each gene is calculated by identifying all the peak regions that intersected with the gene’s body (region between gene start and end site; thus, including introns and UTR regions) a sum of cut-sites across all the intersected peaks is used to ascribe a score to the gene. Gene scores were used as proxy to estimate the potential transcriptional activity of each gene.

##### Poor quality cell filtering

A cluster of cells which had high gene scores for BCL2 and NOX4 genes which are markers for apoptotic cells was identified. Interestingly, these cells also displayed a lower number of accessible peaks in the genes but not overall fewer accessible peaks. These cells were lost when a conservative threshold to the number of accessible genes per cell (> 7500 and < 25000) was set. Peaks and genes are considered accessible if they have a non-zero number of cut-sites. Additionally, a cluster of cells that had a high (>80%) representation from only one of the replicate samples was removed.

##### Proximal and distal peak identification

Cellranger’s gene annotation (‘refdata-cellranger-GRCh38-3.0.0’) was used to mark the TSS region of transcripts. The −2000 to +500 TSS region of genes was considered as promoters. Peaks that overlapped with promoter regions were designated as ‘proximal peaks’. Peaks that didn’t overlap with either the promoter regions or gene bodies were considered ‘distal peaks’.

##### Enhancer matrix generation

The distal peak-set were intersected with enhancer coordinates from GeneHancer database[31] using pyBedTools[47] using default parameters to create an enhancer peak-set. The enhancer peak accessibility was calculated by summing the TF-IDF normalized values of all peaks within each enhancer. For TF-IDF normalization, inverse document frequency was calculated based on all the peaks, but term frequency was calculated only for distal peaks. This matrix was rescaled by dividing values for each cell by the sum of values in that cell and multiplying by 1000.

##### Motif identification and motif matrix generation

The nucleotide sequence for each peak was fetched and saved in FASTA format. The position frequency matrices (PFM) of 579 transcription factor binding sites were downloaded from the JASPAR database (‘JASPAR2018_CORE_vertebrates_non-redundant_pfms_meme’). The individual PFM were loaded into FIMO (MEME Suite 4.12.0) along with the FASTA file to identify significant occurrences of TFBS motifs across the peaks. Only those TFBS motifs occurrences were kept that had p-value < 0.05 and whose score was larger than a threshold value for that motif. The threshold value was calculated by taking the difference between the maximum and minimum score for that motif and multiplying it by 0.25. If multiple occurrences of the same motif were detected in a peak, then the occurrence with the maximum score was considered.

For each of the 579 motifs, their occurrence sites were split into distal and proximal categories based on the corresponding category of the peaks where they were found. Distal and proximal motif-cell matrices were generated for yBM CD34+ cells by summing the TF-IDF normalized value (as done previously for enhancers) of each motif’s corresponding peaks.

#### UMAP and clustering

##### yBM CD34+ cells

The cell-peak (all peaks, except those filtered as mentioned in the ‘Peak filtering’ section) matrices were processed using Scarf (version 0.4.4). Top 100,000 highly variable peaks which were accessible (non-zero cut-sites) in at least 100 cells were identified using the ‘mark_hvgs’ function in Scarf. These values (number cut-sites) in these 100K peaks were library scaled normalized (log transformation was disabled) and used to perform PCA reduction (PCA was trained using only BM12 replicate to minimize batch effects). Top 51 principal components were used to create a KNN graph of cells (15 neighbours). The KNN graph was used to calculate UMAP (min_dist=1, spread=2, n_epochs=2000) embedding of cells. The KNN graph was then used to perform Leiden clustering with resolution parameter set at 0.5.

##### yBM CD38-cells

The yBM cells were filtered based on nCounts (between 20000 and 175000) and nFeatures (between 10000 and 50000), additionally ENCODE blacklisted regions and autosomes were excluded as done previously for yBM CD34+ cells. Top 50,000 most variable peaks which were accessible in at least 50 cells were used. These 50K peaks only included those peaks which had log2 mean normalized expression less than 1 (mean was calculated using only those cells where the peak had non-zero value). The KNN graph of cells was constructed using 51 principal components (PCA trained only on BM12 sample) and using 11 nearest neighbours. UMAP embedding was generated with the same parameters as used for yBM CD34+ cells. Leiden clustering of cells was performed with the resolution parameter set at 0.5.

#### Pseudotime analysis of yBM CD34+ cells (scATAC-seq)

Trajectory analysis was performed using Slingshot. The MTX files containing cell-peak matrix were loaded into Seurat and normalized using Seurat’s Normalize data function. The Seurat object was converted into SingleCellExperiment object and only the filtered cells and 100K most variable peaks (identified by Scarf above) were retained in the object. The cluster membership of cells and UMAP embedding (as calculated by Scarf) was imported into the object. The object was fed into Slingshot with parameters: *‘clusterLabels = ‘scarf_clusts’, extend=‘n’, reducedDim = ‘UMAP’, start*.*clus=8’* wherein 8 denotes the HSC cluster.

#### Identification of enhancer clusters

To create the enhancer clusters for each lineage trajectory, the cell-enhancer peak was subsetted for the cells that had a Slingshot predicted weight value of 0.75 for that trajectory. The cells in this subsetted matrix were ordered based on the pseudotime values of the cells and rolling mean transformation was applied to each enhancer with a window size of 200. Thereafter, standard scaling transformation was applied to the enhancer. Thereafter, for each enhancer, the cells were grouped into 100 evenly sized bins along the pseudotime and a mean value was calculated for each bin. The resulting smoothened and binned matrix was subjected to hierarchical clustering using correlation metric as distance function and ward method for calculating linkage. The dendrogram obtained was subjected to a straight cut with the aim of obtaining 20 clusters.

#### Enrichment of motifs in enhancers

Test of enrichment of TFBS motifs in enhancer clusters was performed using Fisher’s exact test (scipy.stats.fisher_exact). The test was performed individually on each of the six lineage trajectories. Enhancers clusters from each trajectory were grouped into categories: HSC enhancers and terminal enhancers (the grouping scheme is shown in the TableS 5). For each category, the hypothesis of whether the enhancers in the category had a higher occurrence of a given TFBS motif compared to the rest of the enhancers was tested. For the HSC enhancers category, the motifs that had a p-value < 1e-4 in all six trajectories were identified.

#### Establishing enhancer-linked genes

Target genes for enhancer were obtained from GeneHancer database. Only those targets that had an interaction score higher than 5 were considered. Additionally, for strict HSC enhancers only those target genes that had at least two enhancer-target links were considered. The expression of enhancer target genes was queried in the scRNA-Seq data. The gene expression values were smoothened and imputed using MAGIC algorithm[48]. MAGIC (magic-impute Python package) was used with the default parameters. MAGIC imputed values were also used for visualization of expression of TF genes for the corresponding TFBS motifs. Expression values of only those TFBS motifs were queried in scRNA-Seq which were either present in all six trajectories in the case of HSC enhancers or were present enriched in terminal enhancers with p-value < 1e-10.

#### Projection of CD34, CD38, CD11A and CD35 populations

Projection of CD34+/CD38-cell populations of yBM and aBM onto yBM CD34+ or CD38-cells was done using Scarf. Same methodology (for projection and predicting cell clusters) as in the case of scRNA-Seq data was used. Only the top 5 neighbours of each target cell during mapping were recorded.

## Author contributions

G.K. and M.S. conceived and designed the study in consultation with P.D. and D.B.; M.S. designed and performed the experiments with contributions from F.S., R.W., E.E., L.G.U., A.K.C., R.K.T., and C.B.; P.D. designed and performed the bioinformatics- and computational analyses with input from M.S. and G.K.; M.S, P.D, and G.K. analysed and interpreted data; M.S., P.D., and G.K. prepared the figures; G.K. supervised the study; M.S. wrote the manuscript assisted by L.G.U., P.D. and G.K.; All authors reviewed, edited, and approved the final manuscript.

## Acknowledgement

We like to acknowledge Johan Richter, MD PhD, for aged bone marrow samples and Kristin Holmgren for Cord blood sample collection. Additionally, the personnel of Lund Stem Cell Center FACS core Facility, and the Lund University Center for Translational Genomics for their expert assistance in flow cytometry and single-cell genomics, respectively. This work was supported by grants from the Swedish Cancer Society, The Ragnar Söderberg Foundation, the Knut and Alice Wallenberg Foundation, the Swedish Research Council, the Swedish Society for Medical Research, and the Swedish Childhood Cancer Foundation.

## Conflict of interest disclosure

The authors declare no competing interests.

## Supplemental figure legends

**Supplemental Figure 1.**
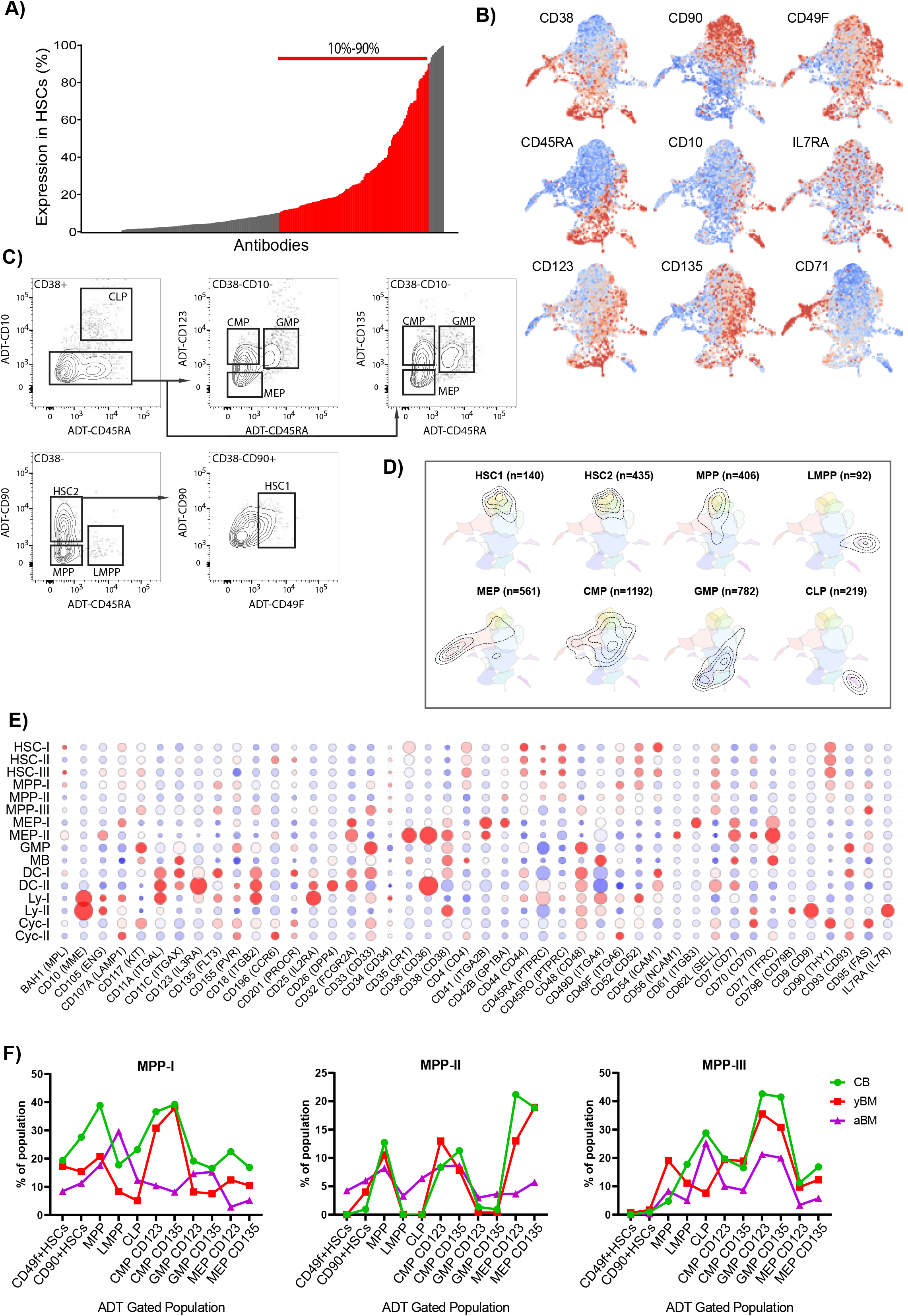
A. Screen of Lin^-^CD34^+^CD38^-^CD90^+^CD45RA^-^CD49f^+^ over 340 unique markers in CB MNCs. B. ADT expression levels of conventional HSPC surface markers on CD34^+^ UMAP, red marks high expression while blue marks low expression. C. Gating of conventional stem and progenitor populations using ADT levels within CD34^+^ cells. D. Gated cells visualized using contour plots on the CD34^+^ UMAP, showing the molecular composition of the gates. E. Bead-plot of expression of surface marker genes within CD34^+^ clusters and the mean ADT level. F. Percentage of molecularly defined RNA-seq subclusters, MPP-I, MPP-II and MPP-III in ADT gated conventional populations in CB, yBM and aBM.

**Supplemental Figure 2.**
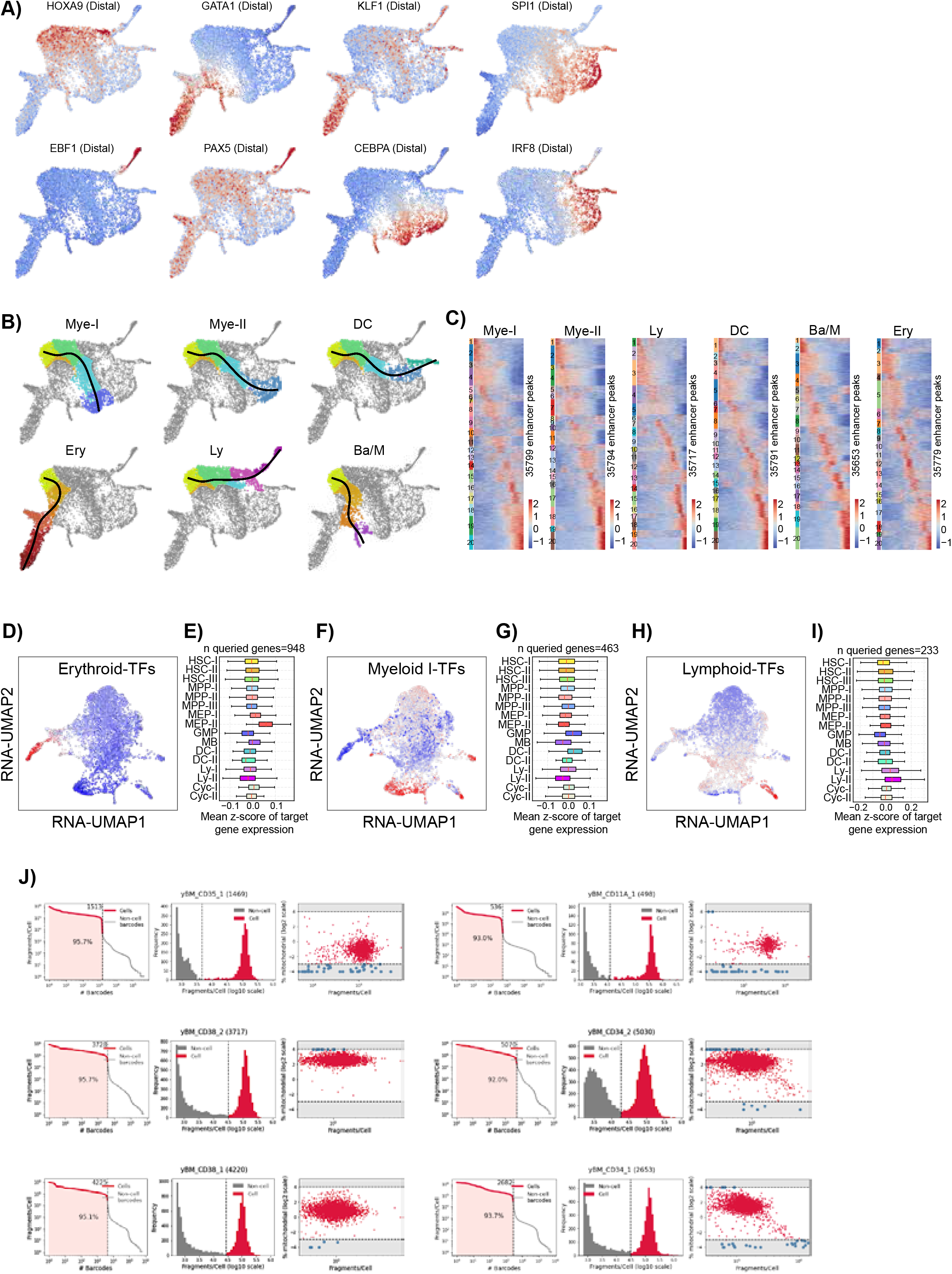
A. Accessibility of key lineage TFBS motifs within the CD34+, red denotes high accessibility and blue low. B. Lineage trajectories for all major lineages defined within the scATAC-seq CD34^+^ HSPCs, all originating from the HSCs. C. Heatmaps of enhancer accessibility across pseudotime of all scATAC-seq trajectories. D, F and H. Cumulative MAGIC mRNA values of terminally enriched enhancer associated transcription factors from Erythroid, Myeloid I and Lymphoid trajectories respectively, displayed on UMAP of yBM CD34^+^ sc-RNA-seq. E, G and I. Mean z-scored mRNA values of enhancer target genes of terminally enriched enhancer from Erythroid, Myeloid I and Lymphoid trajectories respectively, within each cluster of the sc-RNA-seq. J. Filtering and quality control of SC-ATAC-seq samples.

**Supplemental Figure 3.**
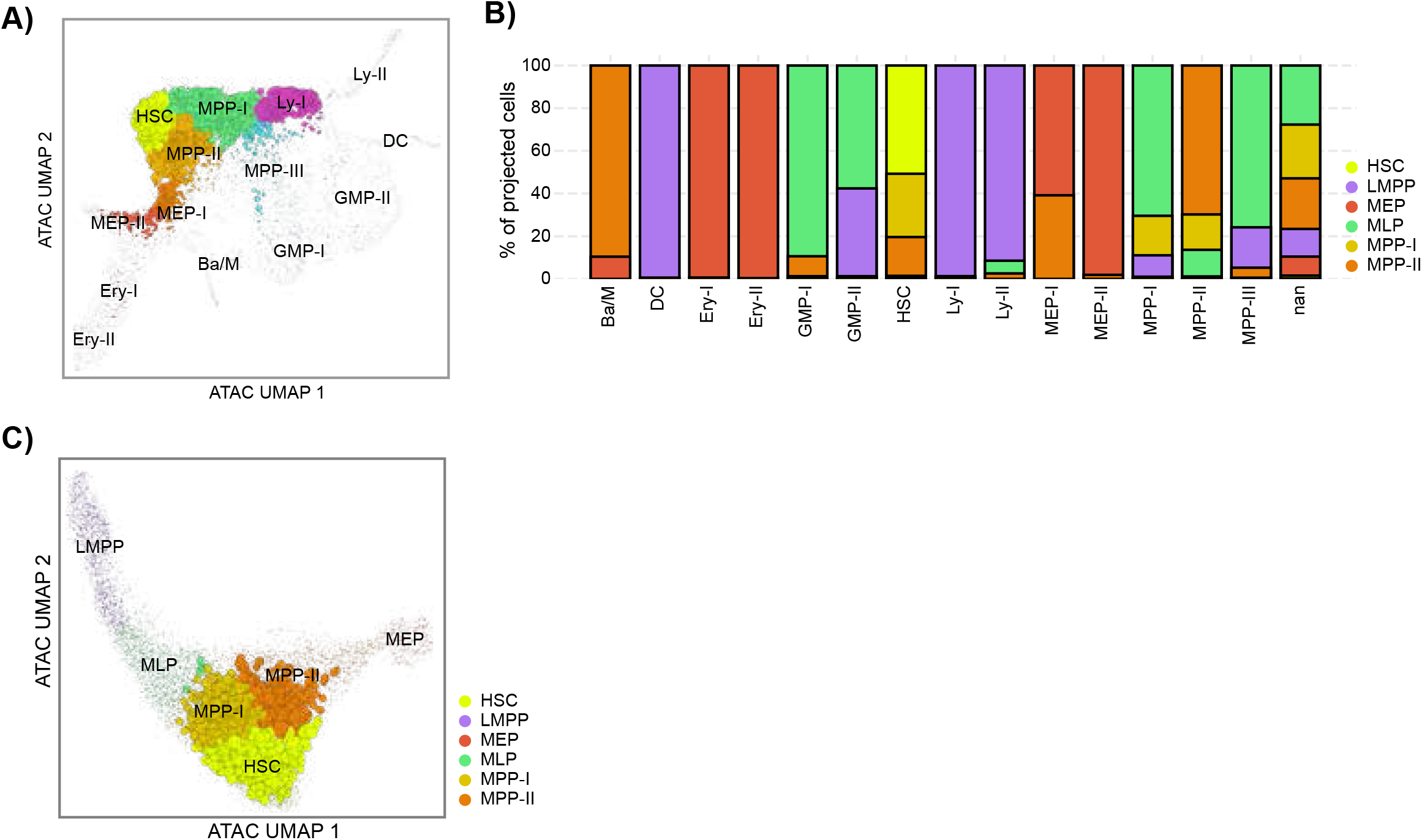
A. SCARF projection of yBM CD38- nuclei onto CD34^+^ scATAC-seq umap. B. Individually projected CD34^+^ scATAC-seq clusters on CD38- clusters to their nearest molecular match. C. Projection of HSC cluster of CD34^+^ scATAC-seq onto CD38- scATAC-seq umap.

**Supplemental Figure 4.**
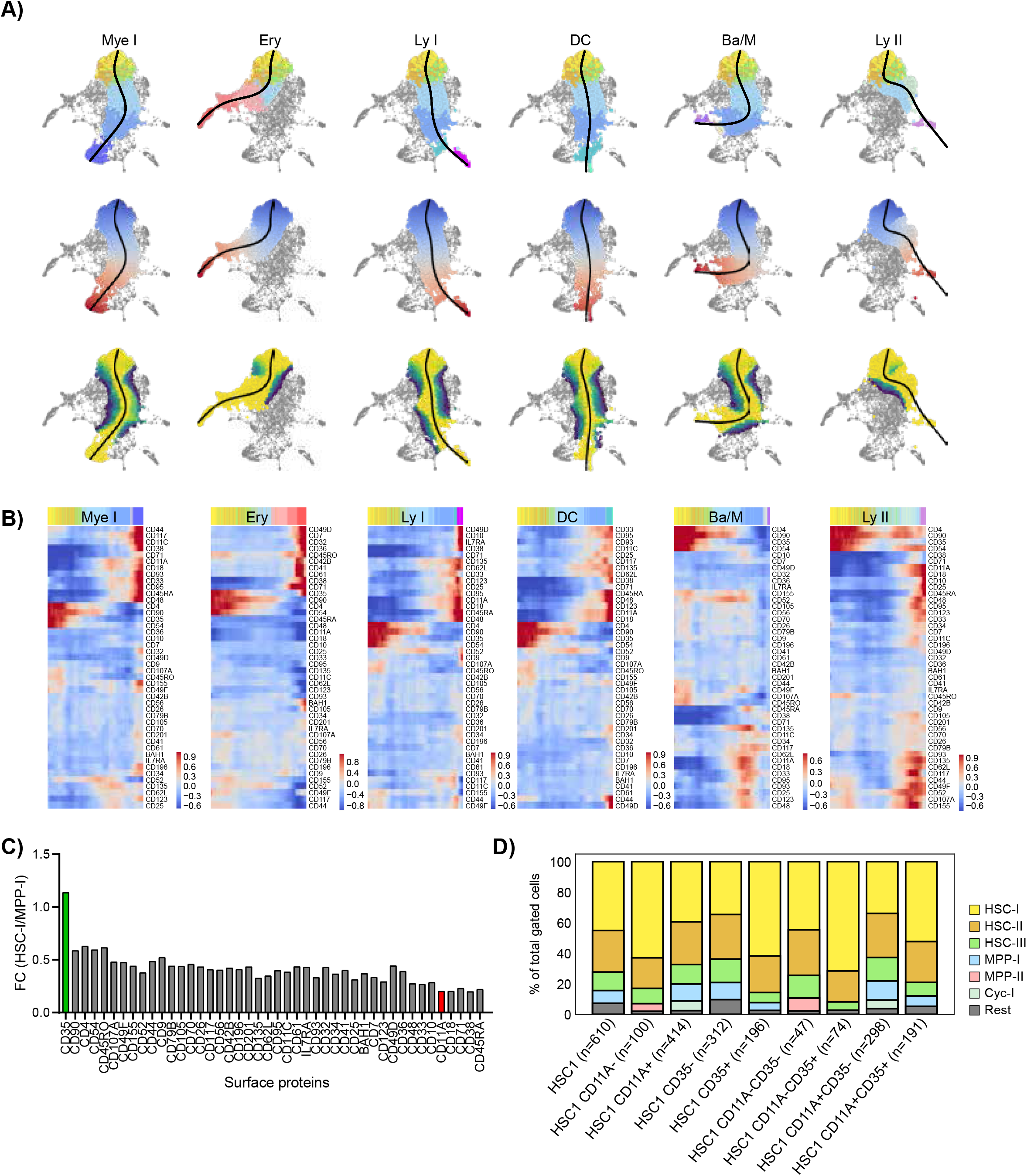
A. Lineage trajectories for all major lineages defined within the CITE-seq CD34^+^ HSPCs, top row; the trajectories along clusters, middle row; pseudotime along trajectories, blue designates early while red designates late pseudotime, and last row confidence of trajectories, yellow designates high confidence, blue designates low confidence. B. Heatmap of ADT expression along pseudotime of all trajectories. C. Fold change of percentage masses of ADTs in HSC-I and MPP-I clusters (% of HSC-I/% of MPP-I). D. ADT-gating of CD90^+^CD49f^+^ (HSC1), CD11A and CD35 within sorted CD38-cells. Proportion of scRNA-seq defined clusters captured within the gated cells.

**Supplemental Figure 5.**
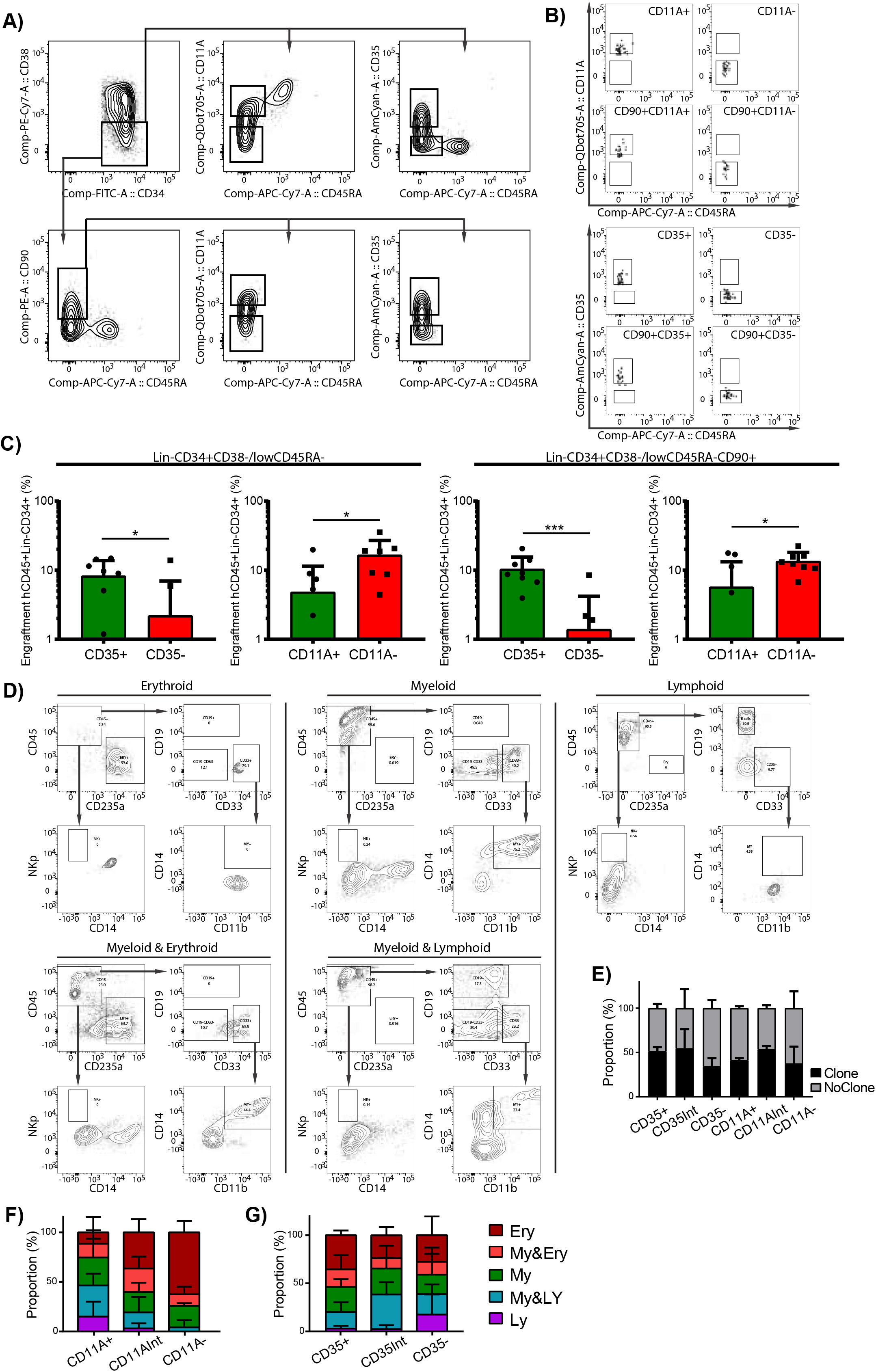
A. Representative image of gateing on CB for transplantation of Lin^-^CD34^+^CD38^-^CD45RA^-^CD35^+/-^, Lin^-^ CD34^+^CD38^-^CD45RA^-^CD11A^+/-^, Lin^-^CD34^+^CD38^-^CD45RA^-^CD90^+^CD35^+/-^ and Lin^-^CD34^+^CD38^-^CD45RA^-^ CD90^+^CD11A^+/-^. B. Reanalysis of sorted cell populations for transplantation assay. C. %CD34^+^ cells in BM after 16W, gated on hCD45^+^Lin^-^(CD19^-^CD33^-^CD15^-^)CD34^+^ cells. D. representative gateing for in-vitro single cell assay. E. Clonal out-put of CD34^+^CD38^-^ sorted single cells within gated CD35^+^, CD35^int^, CD35^-^, CD11A^+^, CD11A^int^ and CD11A^-^. F and G. Lineage output of CD35^+^, CD35^int^, CD35^-^, CD11A^+^, CD11A^int^ and CD11A^-^, dark red represents erythroid, light red is myelo-erythroid, green myeloid, turquoise lympho-myeloid and purple represents lymphoid-output.

## Notes

### Competing Interest Statement

The authors have declared no competing interest.

### Summary of Updates

We found a typo in the title.

